# State-dependent structure-function coupling reflects dynamic engagement of the brain’s structural scaffold

**DOI:** 10.64898/2026.11.12.766152

**Authors:** Robin Pedersen, Roser Sala-Llonch, Jonas Persson, Alireza Salami

## Abstract

Functional brain organization is shaped by the brain’s anatomical scaffold, yet how structure-function coupling varies across cognitive states remains unclear. Across 723 adults from two independent cohorts, we examined coupling using diffusion and functional magnetic resonance imaging. In the primary cohort, coupling was strongest during movie-watching, intermediate during rest, and weakest during working memory, declining further with increasing working-memory load. The movie–rest difference replicated in an independent sample, challenging the primacy of rest for revealing anatomical constraints. State effects were superimposed on a stable unimodal-to-transmodal cortical hierarchy, with movie-watching preferentially enhancing coupling in sensory and language systems. Regions with more stable functional connectivity across states showed stronger coupling. Older age was associated with lower auditory and somatomotor coupling in both cohorts, alongside heterogeneous increases in association cortex. Together, we show that cognitive context shapes structure-function alignment, supporting a framework of complementary structure-constrained and distributed-integrative modes of communication.

## Introduction

Functional brain interactions are dynamic, but they unfold within anatomical constraints imposed by long-range white-matter pathways, local circuits, and polysynaptic network topology (1–3). How anatomical architecture shapes large-scale functional communication is a central question in systems neuroscience, with broad implications for cognition, aging, and brain disorders (3, 4). Structure–function coupling (SFC) quantifies the correspondence between ongoing functional connectivity (FC) and structural connectivity (SC), providing a macroscale index of the extent to which functional interactions follow the brain’s anatomical organization (3, 4). SFC has often been regarded as a relatively stable feature of brain organization. However, accumulating evidence indicates that coupling varies over time (5), differs across individuals (6), relates to cognition (7), and adapts to changes in cognitive context (8). These observations indicate that SFC is not a fixed feature of the structural connectome but varies with the functional interactions expressed across the anatomical scaffold. Yet, why some cognitive states express stronger structure-function alignment than others remains largely unknown.

It is well understood that task engagement reconfigures large-scale functional networks, often shifting functional organization towards more integrated and distributed patterns of interactions (9–15). Tasks differ in the systems they recruit and in the balance they establish in within-system interactions and distributed coordination across systems. Functional interactions can further be supported by multiple complementary mechanisms, including direct anatomical pathways, indirect polysynaptic routes, and distributed interactions spanning different functional systems (3, 16–18). These mechanisms may operate differently across brain regions and cognitive contexts, with their relative contribution depending on the condition and its demands. State-dependent SFC may therefore reflect how a particular functional configuration engages the same structural scaffold.

These shifts occur within an anatomically heterogeneous cortex. Network communication models suggest that the structural features most predictive of FC vary across cortical systems (19). Unimodal sensory regions are organized around specialized processing streams, whereas transmodal association regions have more distributed connectivity spanning multiple systems. This distinction is also apparent in SFC, which is stronger in unimodal cortex and progressively weaker toward transmodal association cortex (20–24). State-dependent differences in SFC may therefore arise both from changes in large-scale functional topology and shifts in the relative engagement of cortical systems that differ in the degree of anatomical constraint.

We propose a framework in which state-dependent SFC can be understood as two complementary modes of structure-aligned interactions. The first is a structure-constrained mode, characterized by concentrated functional interactions that preferentially conform to anatomical pathways. The second is a distributed-integrative mode, in which functional organization increasingly relies on indirect, cross-network, and context-sensitive interactions that are less directly predicted by structural connectivity. These modes should not be regarded as mutually exclusive or as fixed properties of any particular system. Rather, they provide complementary descriptions of how functional interactions may relate to anatomy across regions and cognitive contexts. Consequently, conditions that recruit highly anatomically tethered sensory and multimodal processing streams, such as naturalistic movie-watching, may reflect increased SFC relative to task free conditions. In contrast, conditions that increasingly depend on distributed and integrative coordination across association and control systems, such as a working memory task, may reduce SFC as cognitive demands increase.

This framework generates predictions across multiple levels of brain organization. At a global level, more segregated functional configurations may preserve stronger alignment with brain structure, as concentrated interactions within functional systems may more closely correspond to the structural scaffold (25, 26). However, more integrated configurations may strengthen (27) or weaken (28) the coupling between structure and function depending on the functional configuration. At the regional level, cortical areas with more stable FC profiles across contexts may exhibit stronger SFC, consistent with the possibility that recurrent functional interactions are more closely aligned with anatomical connectivity (20, 29). At an individual level, state-related changes in SFC may depend on how functional connections are reorganized relative to the underlying structural network during task engagement. Age-related differences in SFC may further reflect regional variation in the relationship between structural architecture and functional organization. Previous work has reported greater age-related reductions in coupling in sensorimotor systems, alongside relative preservation and occasional increases in association cortex (19, 30), potentially changing the expression of SFC across cognitive states.

Despite a growing interest in state-dependent SFC, existing evidence of SFC variability between different cognitive contexts remains sparse and fragmented. The few studies comparing resting-state with task-based SFC differ substantially across cognitive paradigms, coupling metrics, global or regional analyses, and methodological approaches (5, 7, 8, 31, 32). Moreover, most studies compare a single task with rest, making it difficult to determine whether observed differences reflect general principles of large-scale communication or task-specific adaptations. Likewise, previous work has rarely examined whether state-dependent changes in SFC emerge coherently across multiple levels of brain organization, including whole-brain coupling, functional network topology, regional cortical organization, individual differences in functional reconfiguration, and aging. A unified framework that links these different organizational levels is therefore still lacking.

Here, we combined diffusion and functional MRI from two adult lifespan cohorts to test how cognitive context shapes structure–function alignment. In the primary cohort from the DyNAMiC study (33), we compared rest, naturalistic movie-watching, and working memory at global, network, regional, and individual levels. A cohort from the Cambridge Centre for Ageing and Neuroscience (CamCAN) study (34, 35) provided an independent test of the movie–rest difference, and extended the comparison to sensorimotor engagement, which allowed us to examine age associations across a broader adult age range. We tested whether functional topology and regional organization account for state-dependent coupling, and whether age associations are shared across states and cohorts. In summary, we found stronger global SFC during movie-watching than rest in both cohorts, whereas working memory showed lower SFC that declined with increasing load. State differences were superimposed on a stable cortical hierarchy, with selective movie-related increases in sensory and language systems and stronger coupling in regions whose FC profiles were more consistent across states. Age associations were regionally heterogeneous, including lower auditory and somatomotor SFC with older age in both cohorts. Together, these findings identify SFC as a dynamic property reflecting how functional organization aligns with the brain’s structural scaffold across cognitive contexts and adulthood.

## Results

### Dataset, sample, and analysis overview

The primary sample from DyNAMiC comprised 177 adults (88 females, 89 males; age range = 20–78 years, mean = 49.89, SD = 17.48), with fMRI during rest (RS; n = 176), movie-watching (MW; n = 174), and a n-back working memory task (WM; n = 175) (33). CamCAN (34, 35) contributed 546 adults (265 females, 281 males; age range = 18.5–88.9 years, mean = 52.53, SD = 17.98), with rest (n = 271), movie-watching (n = 536), and a sensorimotor task (n = 346). Counts for each condition reflects surviving number of scans following high motion exclusions described in Methods. Alongside an independent movie–rest comparison, CamCAN adds a sensorimotor task requiring button presses in response to auditory and/or visual stimuli and broadens the adult age range for examining age associations.

We quantified how closely functional connectivity followed each participant’s structural connectome. Global SFC was computed as the Pearson correlation between structural and functional connection weights across non-zero SC edges, while regional SFC was computed by correlating each region’s structural and functional connectivity profiles. Higher SFC therefore indicates closer correspondence between the relative strengths of structural and functional connections. We then examined whether state-dependent variation in SFC was related to large-scale network topology, regional stability of functional interactions, individual differences in functional reconfiguration and aging. State comparisons accounted for repeated measures, age, sex, and head motion.

### Global SFC is highest during movie-watching

We first tested whether SFC differed at the global level across cognitive contexts in the primary cohort from DyNAMiC. SFC varied significantly across states (F(2, 518) = 47.89, p < 0.001), with covariate-adjusted means showing the strongest coupling during movie-watching (r = 0.250, 95% CI [0.242, 0.259]), intermediate during rest (r = 0.241, 95% CI [0.233, 0.250]), and weakest during working memory (r = 0.228, 95% CI [0.220, 0.237]) (Fig. 1a). Pairwise contrasts confirmed greater coupling during movie-watching than rest (t(518) = 4.04, p < 0.001) and working memory (t(518) = 9.74, p < 0.001), and lower coupling during working memory than rest (t(518) = −5.72, p < 0.001). Within participants, 63.79% of subjects showed MW > RS and 72.41% showed RS > WM, suggesting that group-level state ordering was also evident within individuals (n = 174 for each comparison; Fig. 1b). Full model estimates are reported in Table S1.

**Fig. 1:**
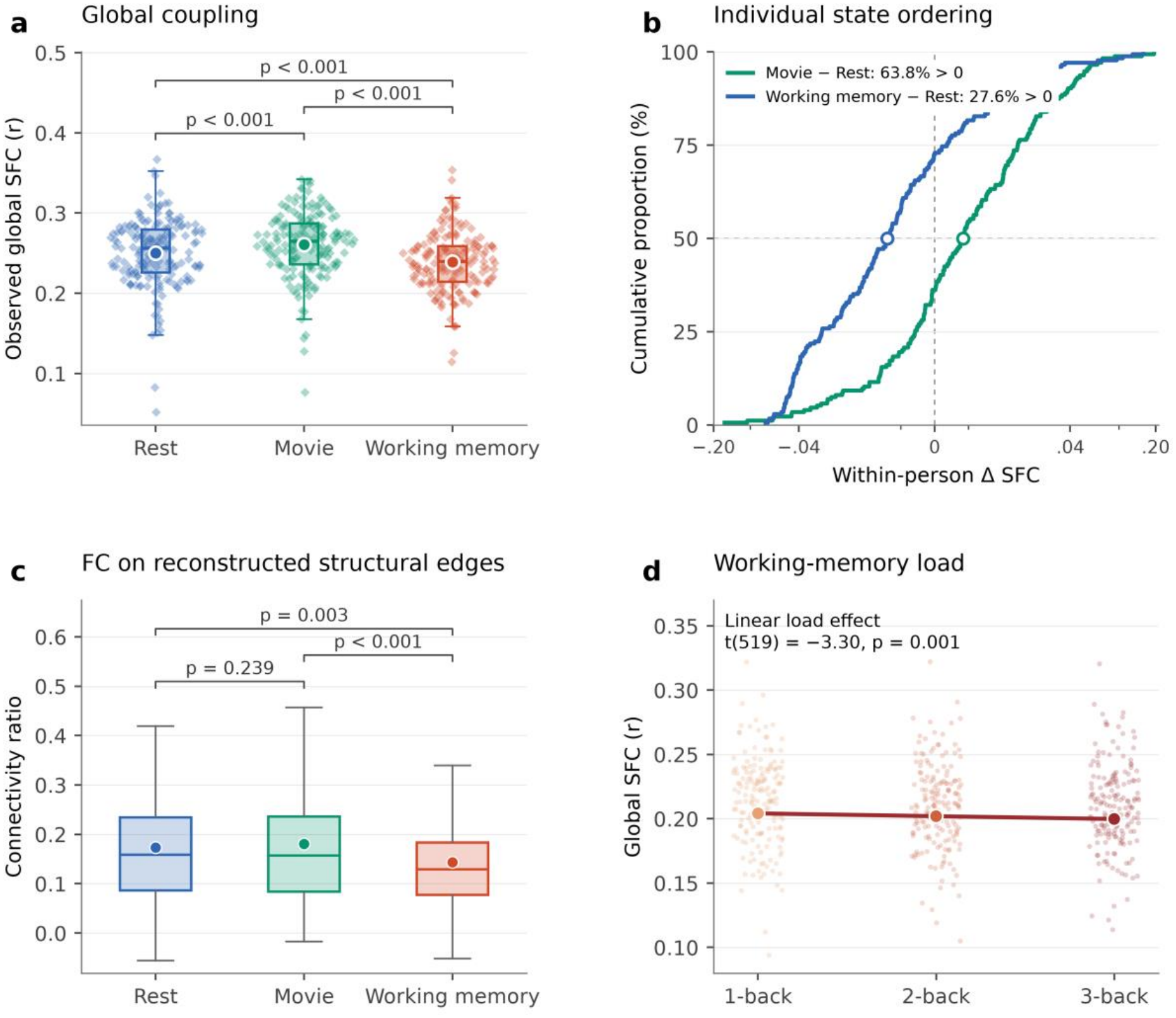
Cognitive state and load shape global SFC. (a) Observed global SFC. N = 176 for rest, 174 for movie-watching, and 175 for working memory. (b) Cumulative distributions of Movie−Rest and Working memory−Rest SFC differences using 174 eligible subjects. The horizontal reference marks 50%; circles mark medians, and the vertical reference marks no difference. (c) Connectivity ratio across conditions. (d) Global SFC across working-memory loads (175 participants). Small points show individual observations; the line and larger points show fitted values from the covariate-adjusted linear load model. Boxes in a and c show medians and interquartile ranges, with whiskers extending to 1.5 interquartile ranges, and circles indicating observed means. Brackets in a and c show two-sided, unadjusted p-values from the covariate-adjusted mixed-model state contrasts (Tables S1 and S2).

We next asked whether state differences in SFC were accompanied by changes in the relative FC strength between structurally interconnected regions versus regions with indirect or weak structural pathways. For each state, we computed a ratio index of FC between regions with non-zero SC weights, relative to pairs of regions without reconstructed white matter streamlines. As such, the index reflects the proportion of FC between regions that are more likely to be directly interconnected rather than sharing indirect or weak pathways. The connectivity ratio was 0.127, 95% CI [0.097, 0.158] during rest, 0.139, 95% CI [0.108, 0.169] during movie-watching, and 0.099, 95% CI [0.069, 0.130] during working memory (Fig. 1c). Movie-watching did not differ significantly from rest (p = 0.239), whereas working memory showed a lower ratio than both rest (t(518) = −3.01, p = 0.003) and movie-watching (t(518) = 4.17, p < 0.001; Table S2). Thus, FC during working memory was more concentrated along structurally weak or indirectly connected regions compared to the resting-state and movie-watching states, converging with the global SFC analysis. These state contrasts were retained after excluding the weakest 25% or 50% structural edges; the working-memory reduction in the connectivity ratio became larger, while the movie–rest ratio remained nonsignificant (Table S3).

Together, these findings show that global SFC is state dependent and characterized by a pattern in which naturalistic movie-watching produced the strongest alignment between functional interactions and the structural scaffold, whereas working memory was associated with weaker alignment and less preferential concentration of FC along direct structural pathways identified by tractography.

### Movie-related SFC increases replicate in an independent sample

We next asked whether the movie-related increase in global SFC was evident in the independent CamCAN sample and whether sensorimotor engagement showed a similar difference from rest. Consistent with our findings in the DyNAMiC cohort, global SFC differed across states (F(2,1147) = 27.90, p < 0.001) with higher SFC during movie-watching than rest (t(1147) = 7.02, p < 0.001). Among 265 participants with both movie and rest runs, 72.83% showed greater SFC during movie-watching, providing within-person support for the replicated state effect (Fig. 2; full model estimates are reported in Table S4). Intriguingly, SFC was also higher during the sensorimotor task than rest (t(1147) = 6.26, p < 0.001), whereas movie-watching and sensorimotor SFC did not significantly differ (p = 0.108).

**Fig. 2:**
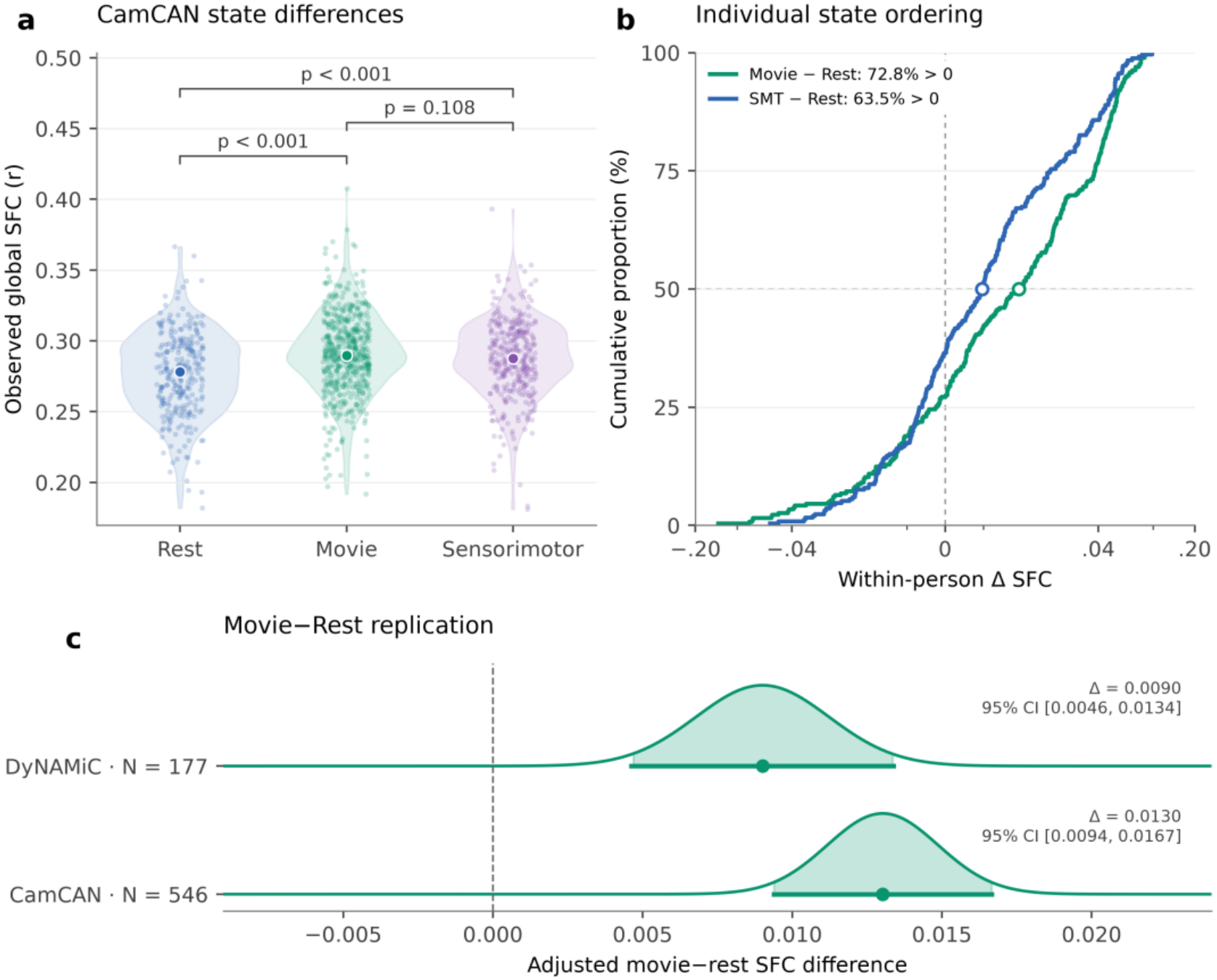
Independent replication of state differences in global SFC. (a) Violin densities and individual observations for eligible runs. Larger dots indicate medians. Rest, movie, and sensorimotor task (SMT) contribute 271, 536, and 346 runs, respectively. (b) Cumulative distributions of paired Movie−Rest and SMT−Rest differences. The horizontal reference marks 50%; circles mark medians, and the vertical reference marks no difference. (c) Adjusted Movie−Rest contrasts in DyNAMiC (177 participants) and CamCAN (546 participants). Curves show t-based confidence distributions centered on model estimates, shaded central regions and horizontal bars indicate 95% confidence intervals (CI). Brackets in a show two-sided, unadjusted p-values from covariate-adjusted mixed-model state contrasts (Table S4).

The independent CamCAN data replicated the increase in global SFC during movie-watching relative to rest and extended this finding to sensorimotor engagement. Together with the reduction in SFC during working memory in DyNAMiC, these findings suggest that the direction of SFC differences relative to rest depends on the cognitive state and may reflect the functional systems recruited. Whether SFC also varies with working-memory load within a given state, however, remains unclear.

### SFC declines with increasing working-memory load

We next examined whether the observed reduction in SFC during the working-memory task represents a general feature of memory engagement or become progressively weaker with increasing working-memory load by comparing coupling across the three levels of the n-back working memory task.

We found that SFC declined from 0.204, 95% CI [0.197, 0.212] during 1-back, to 0.202, 95% CI [0.195, 0.209] during 2-back, and 0.200, 95% CI [0.192, 0.207] during 3-back. A linear load model confirmed that increasing load within the working-memory task was associated with lower SFC (t(519) = −3.30, p = 0.001; n = 175 participants) (Fig. 1d; Table S5).

Thus, SFC decreased progressively with increasing working-memory load even within the same task context. This indicates that reduced SFC during working memory may not solely be attributed to a contextual difference between task conditions but also reflects a systematic shift in the relationship between functional interactions and the structural scaffold as working-memory load increases.

### Functional topology partly explains state-dependent SFC

Because structural connectivity was held constant across states, state-dependent differences in SFC necessarily reflect changes in how functional interactions are organized relative to the structural scaffold. We therefore tested whether state-dependent changes in system segregation, a measure of the relative balance between within- and between-network functional connectivity, could account for variation in SFC across states. As expected, system segregation differed across cognitive states (F(2, 518) = 135.19, p < 0.001). Segregation was highest during rest (adjusted mean = 0.423, 95% CI [0.391, 0.455]), intermediate during movie-watching (adjusted mean = 0.386, 95% CI [0.354, 0.417]), and lowest during working memory (adjusted mean = 0.283, 95% CI [0.251, 0.315]). Both active states therefore shifted functional organization toward a less segregated configuration relative to rest (Fig. 3a; Table S6).

**Fig. 3:**
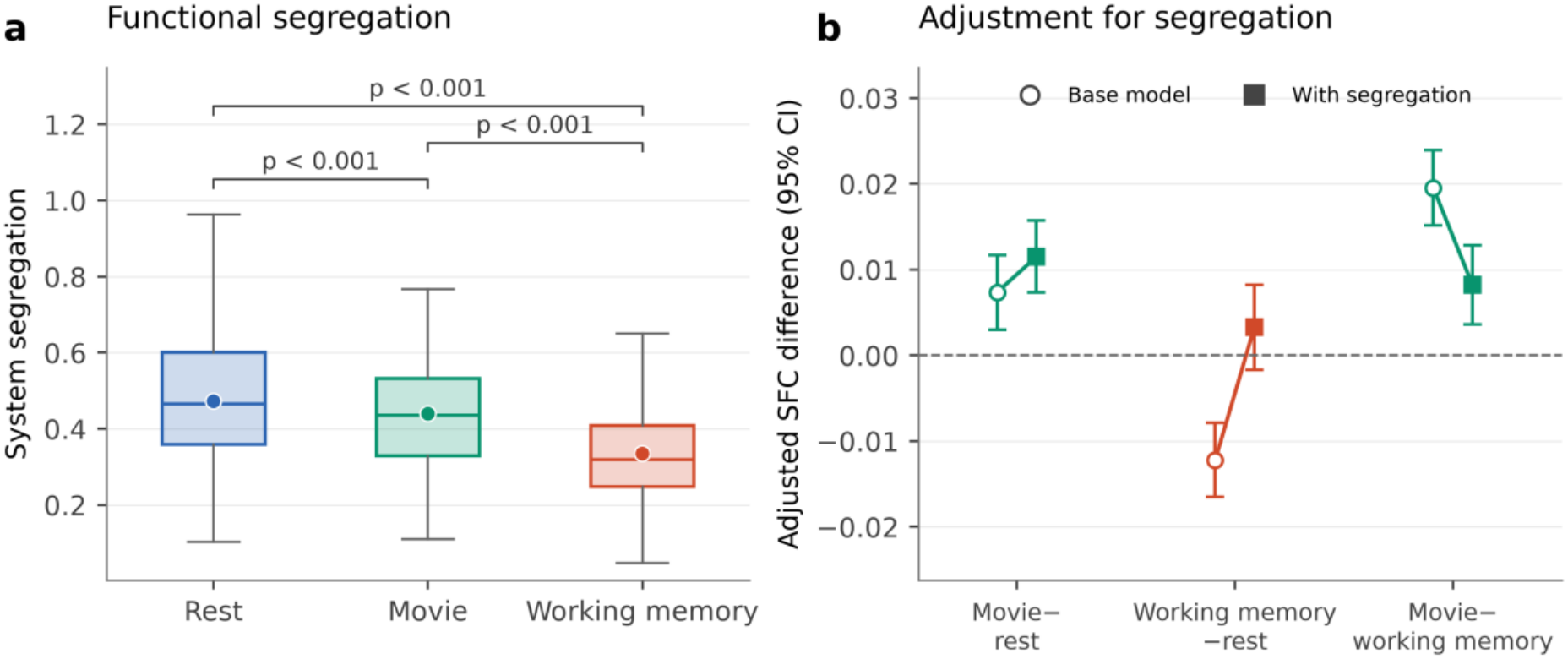
Functional segregation and state differences in SFC. (a) System segregation across states. Boxplots show medians and interquartile ranges, with whiskers extending to 1.5 interquartile ranges, and circles indicating means. (b) Model-adjusted state contrasts and 95% confidence intervals (CI) before and after controlling for state-specific differences in segregation in otherwise identical models (Table S6).

Across all states, segregation was strongly positively associated with SFC (t(517) = 10.97, p < 0.001). Thus, functional interactions were more closely aligned with the structural scaffold when they were less distributed and more concentrated within networks. This association was greater during working memory than rest (t(515) = 3.65, p < 0.001) and movie-watching (t(515) = 2.02, p = 0.044), whereas no difference was observed between rest and movie-watching (p = 0.065). Importantly, when accounting for state-differences in segregation, the magnitude of the observed difference in SFC between rest and working memory was reduced by 72.95%, such that SFC between the two states was no longer significantly different (p = 0.191). However, accounting for segregation differences increased the contrast between movie and rest by 56.52% (adjusted contrast p < 0.001; Fig. 3b; Table S7). Thus, the observed reduction in SFC during working-memory engagement largely reflected a shift toward a less segregated, or more integrated, functional architecture. In contrast, the relative increase in SFC during movie-watching, as compared to rest, could not be explained by network topology alone. This indicates that large-scale functional topology explains much of the reduction in working-memory dependent SFC but does not account for the stronger coupling observed during naturalistic movie-watching. We therefore next asked whether regional patterns of structure–function coupling could explain this remaining difference.

### Movie-watching accentuates the regional SFC hierarchy

Global differences in SFC may arise from spatially heterogeneous changes in how functional interactions engage the structural scaffold. We therefore examined whether state-dependent changes were superimposed on a common regional organization of SFC across the cortical hierarchy. Subject-level regional SFC showed a relatively conserved spatial organization across states (RS–MW: mean r = 0.72, SD = 0.11; RS–WM: mean r = 0.69, SD = 0.11; MW–WM: mean r = 0.67, SD = 0.09).

Across all states, regional SFC followed the canonical unimodal-to-transmodal cortical hierarchy, with the strongest coupling in sensory and somatomotor cortex and the weakest coupling in transmodal association cortex (Fig. 4a). To quantify this organization, we compared the group-level regional SFC pattern with the principal resting-state functional gradient, reflecting differences in functional organization spanning sensory and motor regions to higher-order association cortex (Table S8). Its spatial correspondence with the principal resting-state functional gradient was strongest during movie-watching (Spearman’s ρ = 0.359), followed by working memory (ρ = 0.269) and rest (ρ = 0.256), with greater correlations indicated stronger SFC toward the sensory end of this hierarchy. Bootstrapped state differences supported a stronger correspondence during movie-watching than rest (difference in ρ = 0.103, 95% bootstrap CI [0.082, 0.125]) and working memory (ρ = 0.090, 95% bootstrap CI [0.070, 0.110]). Thus, naturalistic viewing did not reorganize the overall spatial architecture of SFC, but accentuated its existing cortical hierarchy, increasing the differentiation in structural-functional alignment between unimodal and transmodal cortex.

**Fig. 4:**
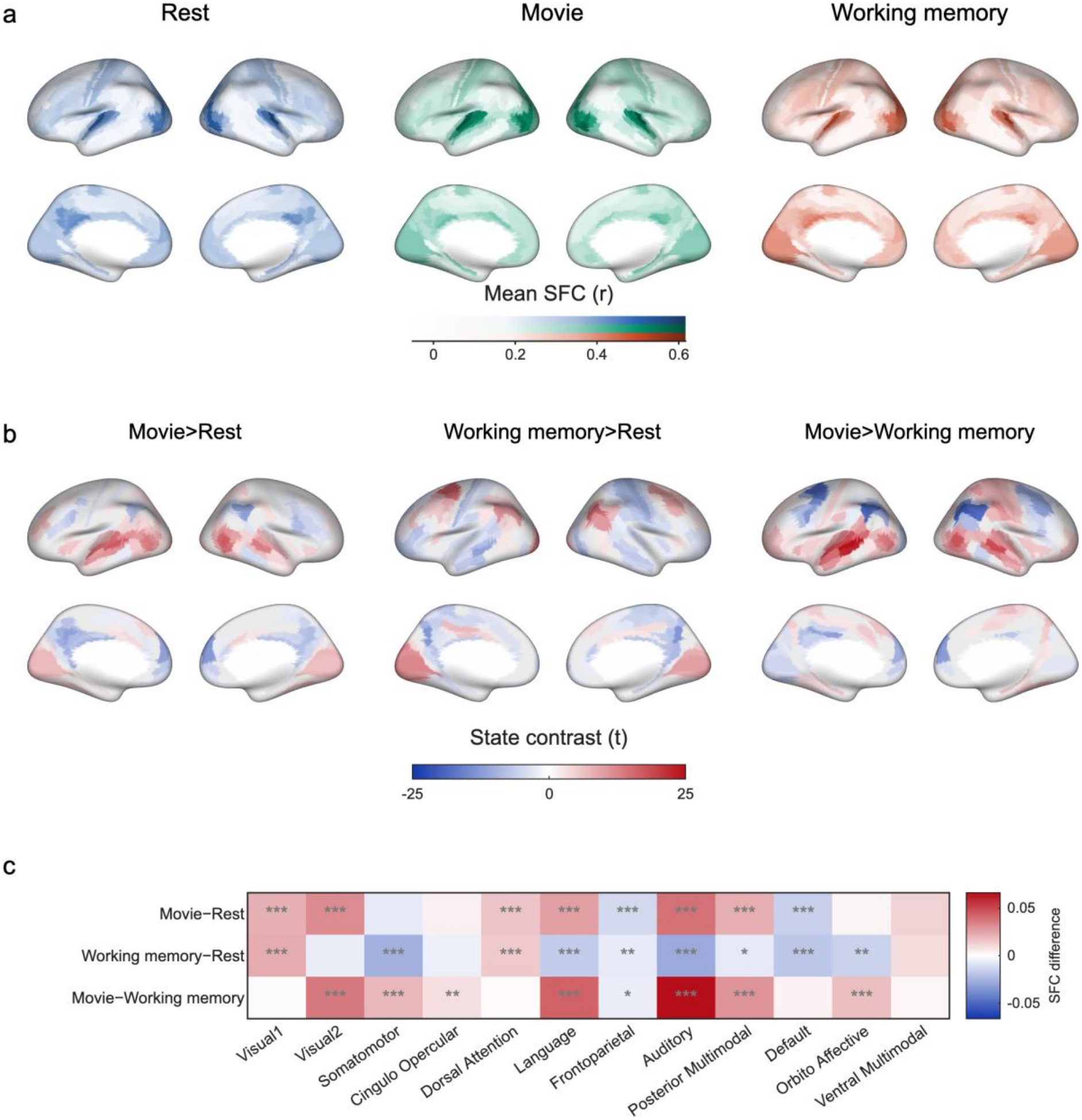
Regional structure-function coupling across states. (a) Mean regional SFC in each state. Color saturation reflects greater SFC. (b) State contrasts from covariate-adjusted regional models; blue–white–red colors indicate t-statistics, with only parcels passing FDR correction (1,080 tests) displayed. Warmer colors indicate stronger coupling while cooler colors indicate weaker relative to the reference condition. (c) Differences in participant-level network means. One, two and three asterisks indicate FDR-adjusted p < 0.05, p < 0.01 and p < 0.001, respectively, across 36 contrasts.

Despite this relatively stable hierarchical backbone, our framework predicts that cognitive states should selectively redistribute structure-function alignment across cortical systems according to their functional demands. We therefore examined regional state contrasts to determine which systems contributed to the observed state-dependent differences in global SFC (Fig. 4b, c). Movie-watching increased coupling in visual, auditory, language, posterior-multimodal, and dorsal-attention systems relative to rest, while default-mode and frontoparietal systems showed lower coupling. Working memory increased coupling in the primary visual and dorsal-attention systems, but showed lower coupling in auditory, somatomotor, language, posterior-multimodal, default-mode, frontoparietal, and orbito-affective systems (Fig. 4c; Table S9). CamCAN similarly showed movie-related increases in visual, language, posterior-multimodal, and dorsal-attention coupling. Notably, the sensorimotor task expressed higher somatomotor coupling together with increases in visual and dorsal-attention systems (Fig. S1; Table S10).

Thus, state-dependent differences in global SFC reflected a combination of regional increases and decreases in structure–function alignment. Movie-watching preferentially increased coupling in sensory and multimodal systems, whereas working memory combined increases in visual and dorsal-attention systems with more widespread reductions elsewhere. Both states nevertheless showed lower coupling in default-mode and frontoparietal systems relative to rest, indicating that distinct global effects can coexist with shared regional responses. The shared movie-related increases in selected sensory and multimodal systems in CamCAN, together with the somatomotor increase during sensorimotor engagement, further support a systematic relationship between cognitive state and the spatial distribution of SFC. These findings suggest that cognitive demands shape the regional pattern of structure–function alignment, with the balance of increases and decreases contributing to the observed global differences.

### Regional functional stability is associated with SFC

Although regional state contrasts exhibited context-dependent deviations, the overall cortical hierarchy was strikingly preserved across cognitive contexts. This raises the possibility that the cortical hierarchy of SFC reflects a more fundamental organizational principle. We hypothesized that regions exhibiting more stable functional interactions across contexts also exhibit stronger alignment with the structural connectome. We therefore examined whether regional SFC was related to the stability of FC across states. For each cortical region, functional stability was quantified from the similarity of its connectivity profile across rest, movie-watching, and working memory, with higher values indicating stronger and more consistent cross-state correspondence. This measure describes similarity across scan conditions, rather than the temporal stability of FC within a scan.

Regions with greater functional stability showed higher SFC in all three states (Fig. 5). Correlations with the group-level stability measure were ρ = 0.372 during rest, ρ = 0.428 during movie-watching, and ρ = 0.370 during working memory (all pspin = 0.002, FDR-adjusted). Bootstrapped state differences supported a stronger correspondence during movie-watching than rest (difference in ρ = 0.056, 95% bootstrap CI [0.041, 0.074]) and working memory (0.058, 95% bootstrap CI [0.042, 0.076]). These findings indicate that regions whose FC profiles were more stable across states showed stronger alignment with SC.

**Fig. 5:**
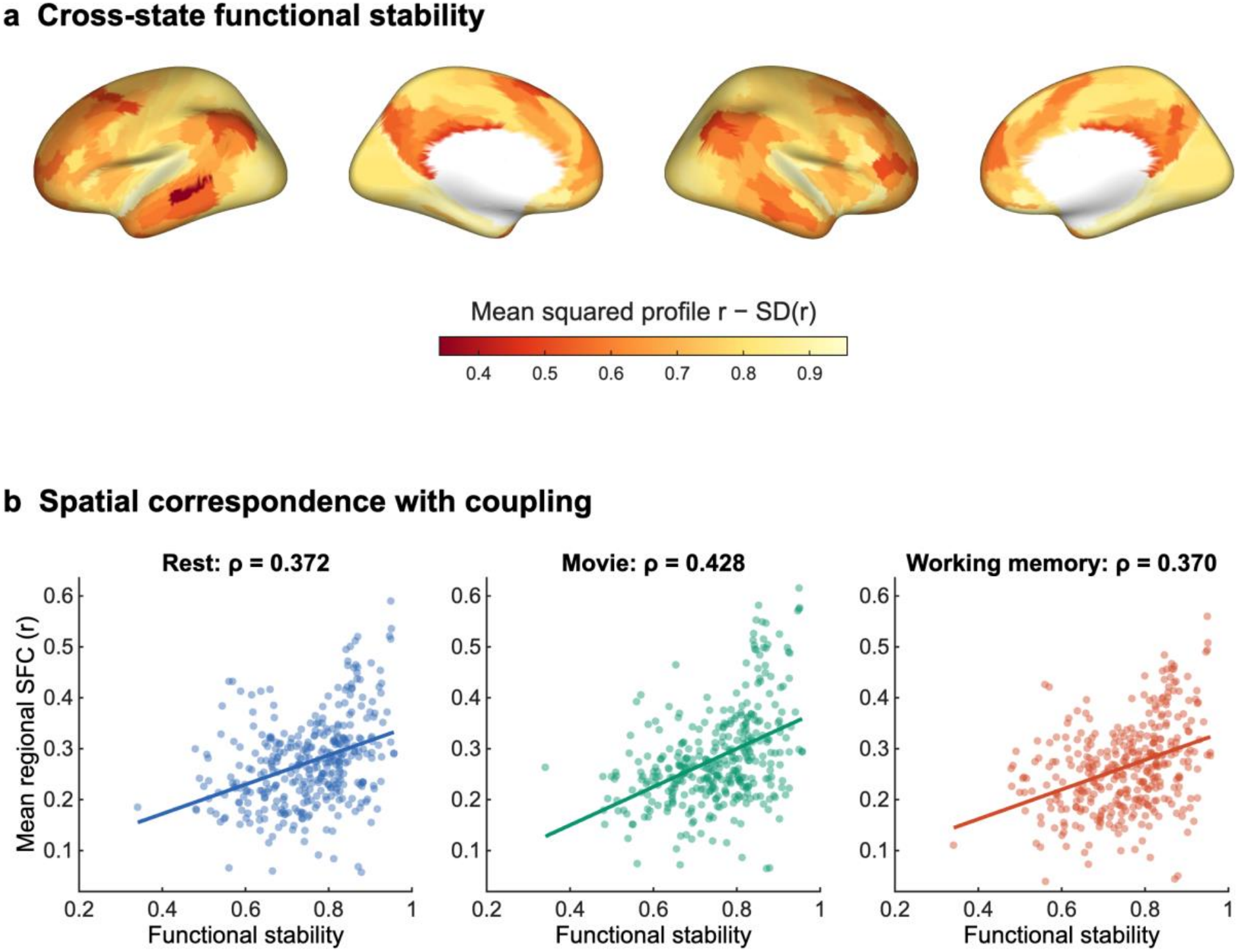
Functional stability and regional structure–function coupling (SFC). (a) Group-level stability, calculated as the mean of the three squared between-state FC-profile correlations minus the standard deviation. Lighter colors indicating greater stability. (b) Correspondence with mean regional SFC in each state. Points represent cortical parcels, lines show linear fits. Reported correlations are Spearman coefficients.

### Functional reconfiguration relates to task–rest SFC changes

The state comparisons describe average differences in SFC, but participants may differ in how their functional connections change during movie-watching and working memory. Greater FC reconfiguration could accompany either stronger or weaker alignment with anatomy; we therefore tested how the extent of reconfiguration related to task–rest differences in SFC. Functional reconfiguration was measured as the dissimilarity between each participant’s resting-state and task FC patterns, with higher values indicating a greater change in the pattern of functional connections. We related this measure to task-minus-rest SFC, such that positive values indicate greater structural–functional alignment during task, whereas negative values indicate greater alignment during rest, and zero represents no difference.

Functional reconfiguration from rest to task was greater during working memory (mean = 0.435, SD = 0.124) than movie-watching (mean = 0.394, SD = 0.123; paired t(171) = 7.51, p < 0.001; Fig. 6a). In a joint model accounting for age, sex, and motion, greater reconfiguration was associated with a more positive task-minus-rest SFC difference during both movie-watching (t(338) = 3.38, p < 0.001) and working memory (t(338) = 4.33, p < 0.001). Thus, participants with greater FC reconfiguration tended to show a larger increase, or a smaller decrease, in SFC from rest to task. Greater reconfiguration therefore did not simply reflect a greater departure from the structural scaffold. However, the associations did not differ significantly between tasks (p = 0.486; Fig. 6b–c; Table S11).

**Fig. 6:**
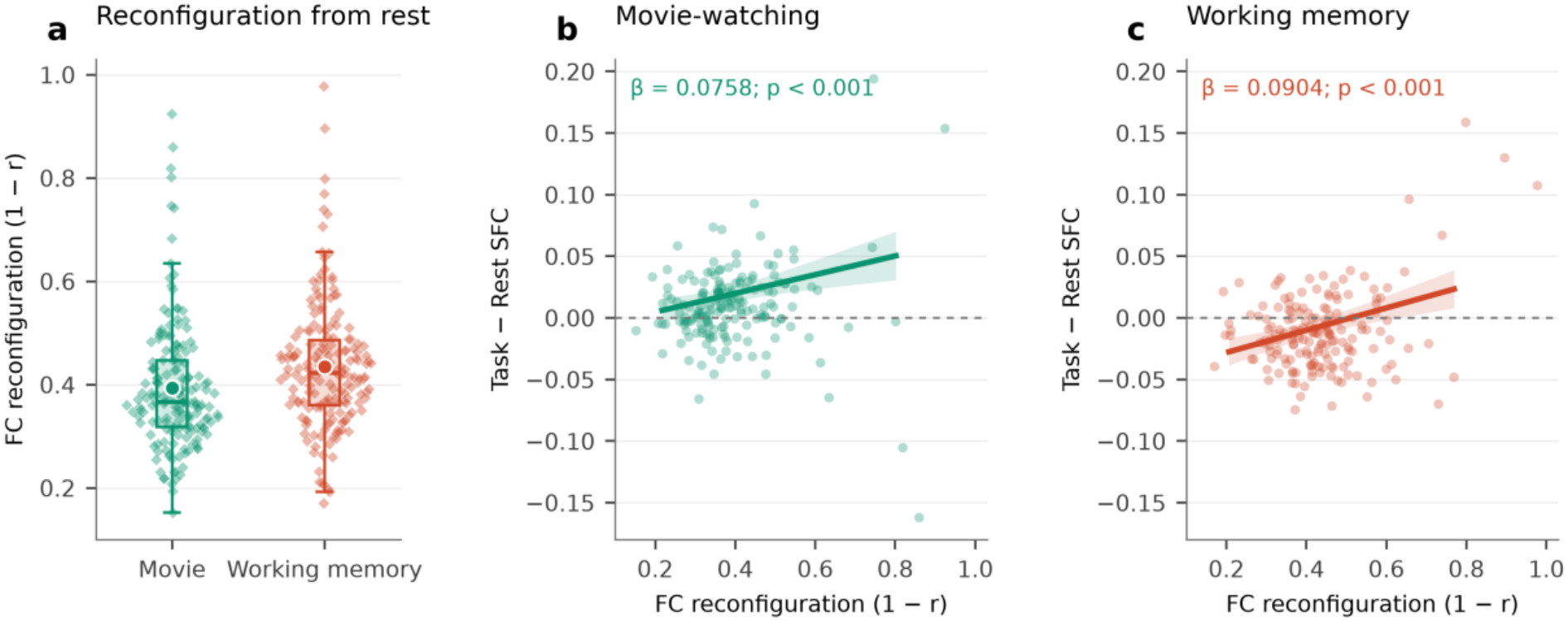
Functional reconfiguration and individual differences in SFC. (a) Functional reconfiguration from rest to movie-watching and working memory, measured as 1 − Pearson r between FC patterns. Points show individual observations, boxes show the median and interquartile range, whiskers extend to 1.5 times the interquartile range, and circles indicate means. Each task–rest comparison includes 174 participants; 172 contribute to the paired comparison between tasks. (b–c) Associations between reconfiguration and task-minus-rest SFC during movie-watching and working memory. Points show observed values; lines and shaded bands show adjusted model predictions and 95% confidence intervals. Positive SFC differences indicate greater coupling during task than rest. Full model estimates are reported in Table S11.

### Age associations with SFC vary across systems and states

If SFC reflects the adaptive engagement of the structural scaffold across cognitive states, age-related differences in functional network organization may alter this capacity. We therefore examined age associations with SFC in both cohorts, focusing on age differences in global SFC, how cognitive states differ from each other, and regional age differences in SFC across the cortex.

Global SFC showed significant age associations across all states in both cohorts, although the shape of the age trajectories differed across states (all padj ≤ 0.016). In DyNAMiC, SFC declined from early to middle adulthood, leveling off in older age during rest and movie-watching, while an apparent late-life upturn was observed during working-memory (Fig. 7a). Age trajectories in CamCAN expressed a similar pattern of decline from early to middle adulthood. However, an upward curvature in older ages was observed for all three conditions, particularly during resting-state (Fig. 7b; Table S12). An overall state-by-age interaction was significant in both cohorts (DyNAMiC: F(4, 514) = 2.39, p = 0.0497; CamCAN: F(4, 1142) = 3.17, p = 0.013). Follow-up comparisons identified a global SFC movie-rest difference in CamCAN (padj = 0.008), although the same comparison did not survive multiple correction in DyNAMiC (Table S13).

**Fig. 7:**
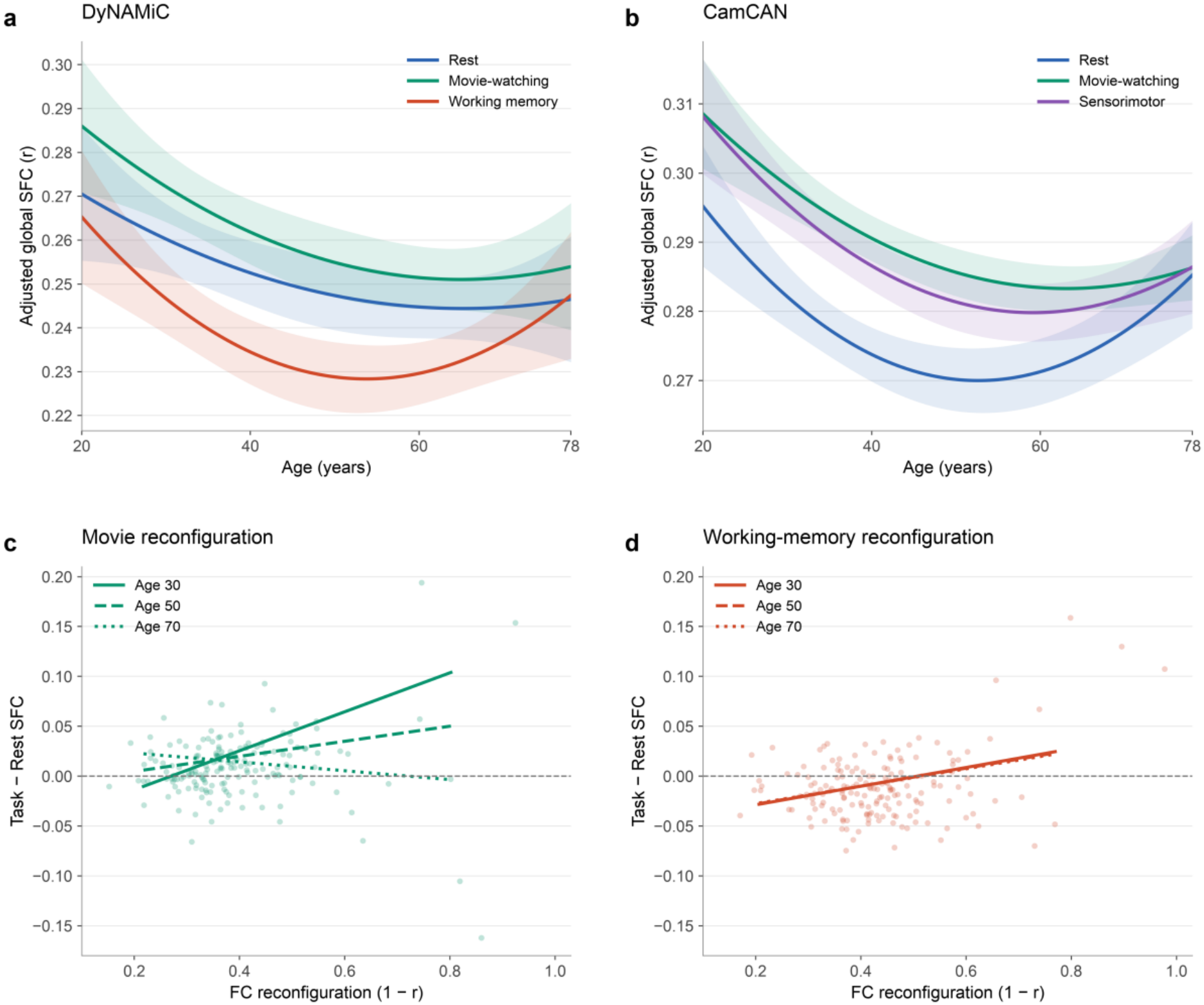
Age associations with SFC and functional reconfiguration. (a–b) Global SFC across age in DyNAMiC and CamCAN. Lines show quadratic mixed-model predictions over the shared 20–78-year age range; shading shows pointwise 95% confidence intervals. (c–d) DyNAMiC task-minus-rest SFC against functional reconfiguration (1 − Pearson r), with fitted relationships at ages 30, 50 and 70 years. Each comparison includes 174 participants. Dashed horizontal lines mark no SFC difference. Overall age and reconfiguration tests are reported in Tables S12 and S11, respectively.

Regional analyses revealed a spatially heterogeneous pattern of age differences (Fig. 8). Across states and cohorts, older age was consistently related with lower SFC in auditory and somatomotor cortex (network summaries in Tables S14 and S15 and Figs. S2, S4 and S5). In DyNAMiC, we primarily observed age-related reductions in coupling in auditory cortex, alongside age-related increases among dorsal-attention and frontoparietal regions. The regional age effects were generally greater in CamCAN, showing a broader mixture of increased coupling, primarily in the dorsal-attention network, alongside reduced coupling in sensory and default-mode regions. While regional age-effects did not differ between states in DyNAMiC, age-related decline in CamCAN was generally greater in somatomotor regions during the two task states when compared to rest (Figs. S2–S3).

**Fig. 8:**
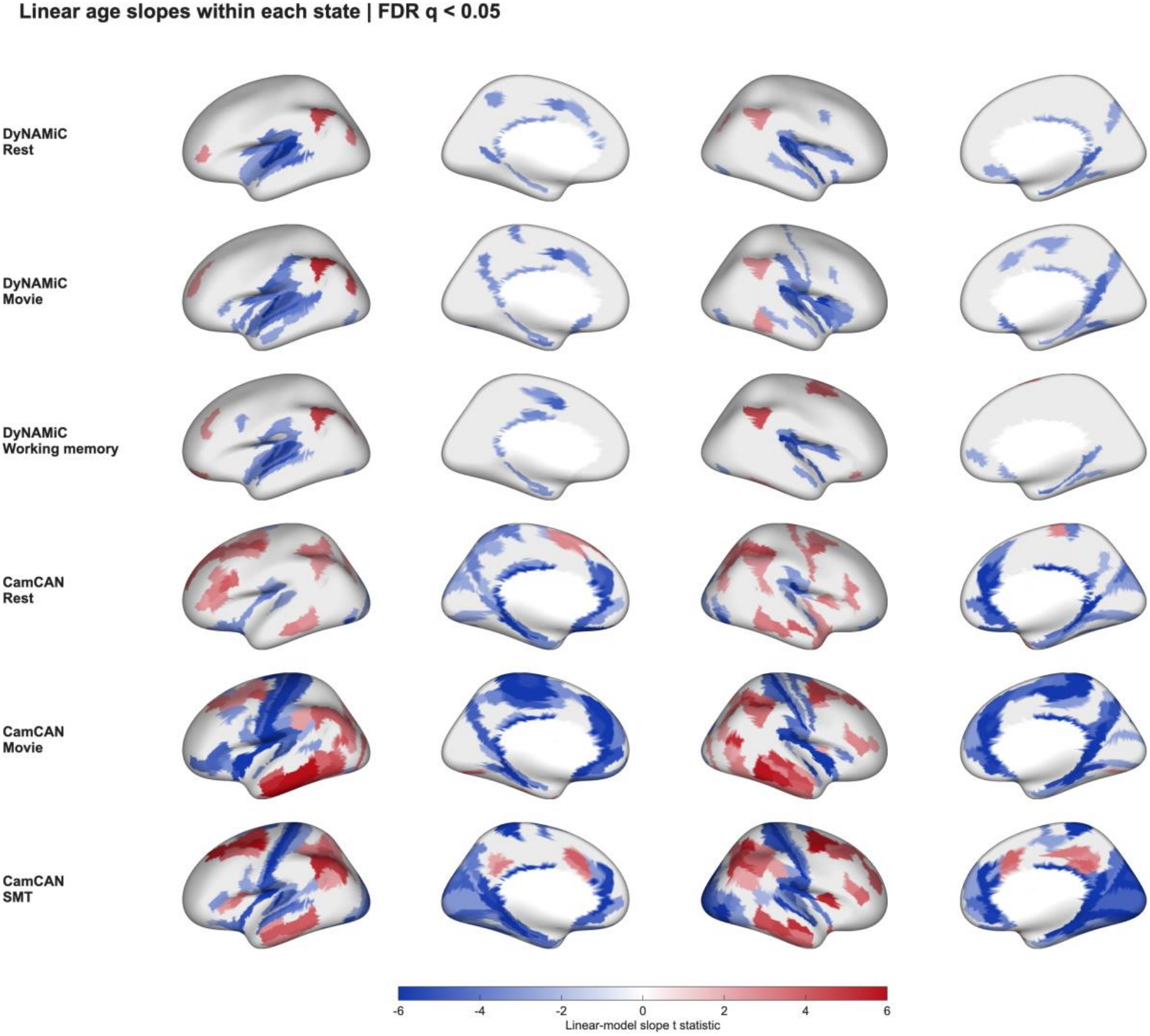
Regional age associations with structure–function coupling. Signed t-statistics for parcelwise linear age slopes in DyNAMiC and CamCAN, shown separately for each state. Blue denotes lower and red higher SFC with older age. Only parcels surviving FDR-adjusted p< 0.05 across 1,080 parcel-by-state tests within each cohort are colored; gray indicates nonsignificant effects; SMT denotes the sensorimotor task.

We next examined whether age moderated the relationship between functional reconfiguration and state-dependent SFC differences. In DyNAMiC, we observed a significant difference in the association between functional reconfiguration and state-dependent difference in SFC across the three states (t(338) = 4.95, p < 0.001; Table S11). Specifically, older age was associated with a weaker relationship between functional reconfiguration and SFC changes during movie-watching relative to rest (t(338) = −5.32, p < 0.001; Fig. 7c), whereas no corresponding age moderation was observed during working memory (p = 0.883; Fig. 7d). Together, these findings indicate that aging is associated not only regional and state-dependent differences in SFC, but also with a context-specific weakening of the relationship between functional reconfiguration and structure-function alignment during movie-watching.

## Discussion

Using within-subject multimodal MRI across 723 adults from two independent cohorts, we demonstrate that SFC is not a fixed property of the connectome but a dynamic systems property that varies systematically with cognitive context. Global SFC was strongest during movie-watching, intermediate during rest, and weakest during working memory. Moreover, coupling declined progressively with increasing working-memory load, indicating that SFC varies parametrically with working-memory load within a given task. Together, these findings suggest that cognitive context shapes the extent to which functional interactions align with the brain’s anatomical scaffold. Specifically, our findings support a conceptual framework involving two complementary modes of large-scale communication. We interpret state-dependent differences in SFC in terms of the relative expression of structure-constrained and distributed-integrative communication, a distinction developed below.

The strongest coupling during movie-watching challenges the widespread assumption that rest provides a privileged context for revealing structure-constrained functional organization. Resting-state FC is highly informative and robustly shaped by structural connectivity, making it the dominant framework for studying structure-function relationships (3, 4). However, rest is also intrinsically unconstrained, allowing spontaneous thought, fluctuations in vigilance, and idiosyncratic internal mentation which introduce substantial variability in ongoing functional interactions (1, 36–39). Naturalistic movie-watching provides a fundamentally different form of cognitive engagement. Continuous, temporally coherent audiovisual, linguistic, social, mnemonic, and predictive structure repeatedly recruits canonical sensory hierarchies while simultaneously engaging distributed higher order systems within a common external temporal structure (40–45). This combination appears to reduce unstructured neural variability while repeatedly engaging functional interactions that remain closely aligned with anatomically supported processing streams. Our findings are consistent with a form of externally structured integration in which large-scale functional communication remains broadly distributed while preserving close alignment with the structural scaffold. By this view, movie-watching occupies a distinct position between unconstrained rest and cognitively demanding task performance, providing an ecologically rich cognitive state in which structure–function alignment was strongest among the three conditions in the two independent samples.

The independent CamCAN analysis replicated the increase in global SFC during movie-watching relative to rest, supporting the generalizability of this state difference across datasets. Higher coupling during the sensorimotor task further indicates that task-related SFC differences depend on the form of cognitive engagement. Importantly, the absence of a significant difference between movie watching and the sensorimotor task suggests that elevated SFC is not unique to naturalistic viewing. Our observation that movie-watching exhibited stronger SFC despite lower functional segregation than rest provides an important insight into the organizational basis of state-dependent SFC. If a reduction in segregation alone were sufficient to reduce SFC, both active states should have shown weaker coupling than rest. Instead, movie-watching increased coupling, whereas working memory decreased it, despite both states shifting the brain toward a more integrative functional configuration. This interpretation is further supported by our finding that accounting for functional segregation reduced the difference in SFC between working memory and rest to a nonsignificant contrast yet accentuated the difference between movie-watching and rest. Together, these observations indicate that functional topology is closely associated with the reduction in SFC during working memory but cannot fully account for the enhanced coupling observed during naturalistic viewing. Thus, the relationship between functional integration and SFC depends not simply on the degree of integration, but also on how functional interactions are organized. These findings support our proposed distinction between structure-constrained and distributed-integrative communication, suggesting that integrated functional communication can differ substantially in its alignment with the structural connectome.

The working memory findings provide complementary evidence for the proposed framework, showing that lower SFC accompanies increasing working-memory load without necessarily indicating less efficient brain organization. Coupling was lower during working memory than during rest and declined across the working-memory loads, indicating that reduced SFC varied parametrically with increasing working-memory load rather than reflecting a simple categorical rest-versus-task difference. Importantly, increasing cognitive demand requires active maintenance, updating, attentional control, and flexible coordination across distributed association and control systems. Such computations are unlikely to be supported exclusively through the dominant anatomical pathways captured by direct structural connectivity. Instead, they may recruit indirect, polysynaptic, and dynamically reconfigurable interactions spanning multiple large-scale networks (10, 12–14). Under this view, reduced SFC during working memory should not be interpreted as a reduction in biological constraint or organizational efficiency. Rather, it is consistent with greater reliance on distributed and context-sensitive functional interactions that are less directly predicted by the structural connectome. Accordingly, our findings suggest that cognitive demands do not simply weaken structure–function coupling but may shift the relative expression of structure-constrained and distributed-integrative communication. This interpretation is consistent with recent evidence that state-dependent SC–FC coupling profiles carry behaviorally relevant information and may reflect cognitive-state-specific communication strategies (7, 8).

The regional analyses further indicated that state-dependent changes in SFC are organized on top of a stable cortical hierarchy rather than through a wholesale reorganization of structure-function relationships. Across all cognitive states, sensory and somatomotor regions showed stronger coupling, whereas association cortex showed weaker coupling, consistent with the well-established unimodal-to-transmodal gradient of structure–function tethering (20–22). We propose that this cortical hierarchy provides the anatomical substrate through which the balance between structure-constrained and distributed-integrative communication is expressed. Regions located toward the unimodal end of the hierarchy are more likely to support communication that remains closely aligned with direct anatomical pathways, whereas transmodal systems provide greater opportunity for flexible, indirect, and context-dependent functional interactions (21–24). Consequently, changes in cognitive state can alter global SFC not only by changing overall network topology but also by shifting the relative contribution of cortical systems that differ in their degree of anatomical constraint. This interpretation is also consistent with emerging evidence that regional variations in SFC reflect multiple converging factors, including myeloarchitecture, cytoarchitecture, neuromodulatory input, intrinsic dynamics, and cortical hierarchy itself (46). Our findings extend this view by adding a state-dependent dimension to this framework, suggesting that cognitive context modulates structure-function alignment within a largely preserved cortical hierarchy, rather than reorganizing the hierarchy itself.

Importantly, different cognitive states did not simply alter the overall magnitude of SFC, they redistributed where structure-constrained communication was most strongly expressed along the cortical hierarchy. Movie-watching preferentially increased coupling in auditory, visual, language-related, and posterior multimodal systems, whereas working memory preferentially enhanced coupling in dorsal-attention cortex and selected parietal and dorsal prefrontal regions while reducing mean coupling in frontoparietal, several sensory, and affective networks. This regional redistribution closely matches the computational demands imposed by each cognitive state. Naturalistic viewing repeatedly engages externally driven sensory, perceptual, and language processing streams whose interactions are strongly supported by recurrent anatomical pathways (40–42, 44, 47). In contrast, working memory requires flexible coordination among distributed association and control systems, where functional interactions increasingly depend on indirect and dynamically reconfigurable communication (10, 12, 14). State-dependent SFC therefore reflects more than a global change in SFC. Rather, cognitive context selectively shifts the cortical systems through which structure-constrained and distributed-integrative communication are expressed.

The strong relationship between regional functional stability and SFC provides a potential explanation for why the cortical hierarchy of SFC is so consistently preserved across cognitive contexts. Regions whose FC profiles remained stable across rest, movie-watching, and working memory also showed stronger regional SFC, suggesting that recurrently expressed functional interactions are more likely to remain aligned with the underlying structural scaffold. This finding supports the idea that high regional SFC reflects more than strong anatomical connectivity alone. Rather, it may identify cortical regions whose patterns of communication are repeatedly expressed across cognitive contexts and therefore remain consistently tethered to anatomical architecture. Conversely, lower regional SFC may characterize regions whose functional interactions are inherently more context-dependent, allowing flexible participation in different large-scale network configurations as task demands change. This interpretation is consistent with previous work showing that association cortex is less tightly tethered to local structural connectivity and participates in more flexible functional coalitions (21–23, 29). More broadly, our findings suggest that regional SFC should not be interpreted as a single measure of better or worse organization. Instead, stronger coupling reflects stable anatomical tethering, whereas weaker coupling may reflect the capacity for flexible, context-sensitive communication.

Our lifespan analyses show that cross-sectional age associations with SFC depend on the spatial level and cognitive state examined. The most consistent network result was lower auditory and somatomotor coupling with older age, including during both rest and movie-watching. This agrees with evidence that age-related reductions in local structure–function correspondence are concentrated in sensorimotor regions, while higher-order systems can show relative preservation (19). The age analyses extend this regional distinction across cognitive states. They also qualify the interpretation of the global age-trajectories, as an overall age association does not by itself establish a specific pattern, and a significant fitted difference between selected ages does not establish a general late-life increase.

Regional increases and decreases can coexist, so global SFC can obscure differences in the anatomical alignment of functional connectivity across cortical systems. Age-related differences in functional segregation and white-matter organization provide plausible contexts for this heterogeneity (48, 49), and previous work has reported increasing structural support for selected regions and network hubs with age (30). Nevertheless, the present findings of age-related increases in dorsal-attention regions did not establish the same age trajectory in both cohorts. CamCAN studies relating structural and functional organization to cognition further indicate that their behavioral significance depends on the network and cognitive process examined (50, 51). Positive slopes or curvature should therefore not be equated with compensation or improved function.

Several limitations of this work should be considered. First, diffusion tractography provides an indirect estimate of anatomical connectivity, and edges without reconstructed streamlines should not be interpreted as true absence of anatomical pathways (52, 53). Second, differences in SFC may partly reflect differences in FC reliability across states. Naturalistic movie-watching often produces reliable stimulus-locked responses across individuals, and higher reliability may increase observed coupling (40, 42, 54, 55). Rest and movie-watching scans had the same duration in DyNAMiC, whereas working memory was slightly shorter (350 versus 330 volumes) and CamCAN used different scan durations and repetition times across states. These acquisition differences and residual differences in data quality may affect FC reliability and observed SFC. Third, state-averaged FC cannot reveal the temporal dynamics through which structure–function alignment changes during specific moments of narrative engagement, attentional orienting, or working-memory demand (5, 56, 57). Fourth, SFC is a standardized correlation, such that an increase can accompany lower SC–FC covariance if FC dispersion decreases more. Greater SFC therefore does not necessarily indicate stronger anatomy-aligned FC amplitude and does not identify the communication mechanisms that generate FC from SC. Future work should therefore combine state-dependent SFC with network communication models that explicitly estimate shortest paths, diffusion, communicability, navigation, controllability, or other indirect routing mechanisms (3, 16–18). Despite these limitations, the consistency of our findings across global, network, regional, individual, and lifespan analyses suggest that the proposed framework captures a robust organizational principle of state-dependent structural-functional coupling. The degree of transmodal structure–function correspondence may also depend on the structural communication model used, as illustrated by recent work indicating improved predictions incorporating high-frequency structural eigenmodes (58).

Taken together, our findings support a reinterpretation of structure-function coupling as a dynamic systems property rather than a static feature of the structural connectome. We propose that the degree of SFC reflects the balance between two complementary modes of large-scale communication; structure-constrained mode and a distributed-integrative mode. In this framework, these modes are proposed to coexist across cognitive states, with their relative contribution varying according to behavioral demands. From this perspective, SFC should not be interpreted simply as a measure of anatomical constraint, nor as an indicator of more or less efficient brain organization. Instead, it provides a systems-level index of how functional communication recruits the brain’s structural scaffold to support different forms of behavior. Although demonstrated here using rest, naturalistic movie-watching, and working memory, in addition to a complementary sensorimotor task, we anticipate that this framework extends more broadly to diverse cognitive states in which the balance between structure-constrained and distributed-integrative communication is continuously adjusted to meet changing behavioral demands. By shifting the question from whether functional interactions are constrained by anatomy to how the brain dynamically recruits its anatomical scaffold to support different forms of behavior, our findings provide a unifying framework for interpreting state-dependent structure–function coupling.

## Materials and Methods

### Participants

#### DyNAMiC

The current study used baseline data from DyNAMiC (33), a prospective study of healthy individuals across the adult lifespan. All participants provided written informed consent. The study was approved by the Regional Ethical Review Board in Umeå, Sweden (approval number 2017/248-31). The study’s design, imaging protocols, and procedures are described in detail elsewhere (33). Here, only methodological and material aspects of relevance for the current study are presented.

The DyNAMiC study participants (N = 180) were recruited by random selection from the population registry of Umeå, Sweden, stratified by six age cohorts between 20 and 80 years (n = 30 per decade, 50% females). Participants were screened for a set of exclusion criteria, including contraindications to magnetic resonance imaging, neurological disorders, brain pathology, cognitive impairment, and medical conditions or treatments potentially affecting brain function and cognition. All participants completed a full set of functional and structural MRI scans. Two subjects were excluded from the current study because of technical issues during tractogram creation. From the remaining 178 participants, runs with mean FD > 0.5 mm were excluded: two rest, four movie, and three working-memory runs, leaving 177 participants (88 females) aged 20–78 years (mean = 49.89, SD = 17.48).

#### CamCAN

The replication used data from the Cambridge Centre for Ageing and Neuroscience (CamCAN) (34, 35). From an initial sample of 635 participants with structural connectomes, resting-state and movie-watching fMRI, three participants lacked sensorimotor-task fMRI, leaving 632 participants before exclusions. We excluded runs with mean FD > 0.5 mm or with more than 50% of volumes flagged for motion-spike regression (framewise FD > 0.2 mm). The latter criterion required at least 131 of 261 non-spike volumes in rest and the sensorimotor task, and at least 97 of 193 in movie-watching. The final analysis included 546 participants with at least one eligible run (250 with all three states), contributing 1153 runs. Other eligible runs from each participant were retained. CamCAN received approval from the Cambridgeshire 2 Research Ethics Committee (10/H0308/50), and participants provided informed consent.

### MRI acquisition

#### DyNAMiC

All MRI data were acquired on a 3T Discovery MR750 scanner (General Electric) equipped with a 32-channel phased-array head coil. High-resolution T1-weighted images were collected using a 3D fast spoiled gradient-echo sequence (176 sagittal slices, 1 mm slice thickness, repetition time [TR] = 8.2 ms, echo time [TE] = 3.2 ms, flip angle = 12^◦^, field of view [FOV] = 250 × 250 mm). Diffusion-weighted images were acquired using a multiband echo-planar imaging sequence with 90 diffusion directions (63 slices, TR = 3000 ms, TE = 73.0 ms, flip angle = 90^◦^, FOV = 256 × 256 mm, b = 2000 s/mm^2^, multiband factor = 3, in-plane acceleration factor = 2). In addition, two sets of 10 b = 0 images with reversed phase-encoding polarity were acquired to enable correction of susceptibility-induced distortions. Whole-brain functional MRI data were acquired during three conditions: resting-state, movie-watching, and working memory using a T2*-weighted single-shot echo-planar imaging sequence with 37 transaxial slices, slice thickness = 3.4 mm, 0.5 mm spacing, TR = 2000 ms, TE = 30 ms, flip angle = 80^◦^, and FOV = 250 × 250 mm. Resting-state and movie-watching scans each comprised 350 volumes acquired over approximately 12 min, whereas the working-memory scan comprised 330 volumes acquired over approximately 11.3 min. All three conditions used the same TR and other acquisition parameters. Ten dummy scans were acquired and discarded at the beginning of each sequence. During resting-state fMRI, participants were instructed to remain awake, keep their eyes open, and fixate on a white cross presented on a black background. During the naturalistic movie-watching scan, participants viewed and listened to a 12-minute video of chronologically ordered sections from the Swedish film “Cockpit” (2012). During working memory, participants performed a numerical n-back task with 1-, 2-, and 3-back conditions (33). Each load comprised nine blocks of ten digits, with blocks presented in random order and preceded by a cue indicating the upcoming load. Digits from 1 to 9 appeared for 1.5 s, followed by a fixation cross for 0.5 s. Participants indicated whether each digit matched the one presented n positions earlier using an MRI-compatible response device, pressing with the right index finger for matches and the right middle finger for nonmatches. Participants completed at least two practice trials before scanning, with additional practice as needed.

#### CamCAN

MRI data was acquired using a 3T Siemens TIM Trio scanner. Resting-state fMRI used a T2*-weighted gradient-echo EPI sequence comprising 261 volumes of 32 axial slices, slice thickness = 3.7 mm, interslice gap = 0.74 mm, TR = 1970 ms, TE = 30 ms, flip angle = 78°, FOV = 192 × 192 mm, in-plane resolution = 3 × 3 mm; acquisition time approximately 8 min 40 s (35). Participants were instructed to lie still with their eyes closed and remain awake (34). During movie-watching, participants viewed an approximately eight-minute version of Alfred Hitchcock’s “Bang! You’re Dead.” The sensorimotor task required button presses in response to auditory and/or visual stimuli (34). The available parcel time series contained 261 volumes at TR = 1.97 s for rest and the sensorimotor task, and 193 volumes at TR = 2.47 s for movie-watching.

### Diffusion MRI preprocessing and structural connectivity

Diffusion-weighted MRI preprocessing and tractography were primarily performed in MRtrix3 (59). Diffusion data were denoised using MP-PCA (60, 61), corrected for Gibbs-ringing artifacts (62), susceptibility-induced distortions, eddy currents, and head motion using FSL’s topup and eddy tools (63, 64). Bias-field correction was applied using the ANTs N4 algorithm (65), followed by brain mask estimation. Tissue-specific response functions for white matter, gray matter, and cerebrospinal fluid were estimated using the Dhollander approach (66, 67).

White-matter fiber orientation distributions were reconstructed using single-shell 3-tissue constrained spherical deconvolution (68), implemented in MRtrix3Tissue (https://3Tissue.github.io), a fork of MRtrix3 (59). The resulting tissue components were intensity-normalized prior to tractography. Whole-brain anatomically constrained tractography was performed using a five-tissue-type image derived from each participant’s T1-weighted scan, which was rigidly registered to diffusion space, with gray matter–white matter interface seeding and iFOD2-based streamline propagation (69, 70). Ten million streamlines were generated, and streamline weights were estimated using SIFT2 (71). For connectome generation, the HCP-MMP1 atlas (72) was projected from fsaverage to individual cortical surfaces using FreeSurfer (73) and converted to a volumetric parcellation in native diffusion space to define network nodes.

Structural connectomes were then constructed from the full tractogram with edges defined as the sum of SIFT2 streamline weights between node pairs, scaled to account for differences in node volume (74). Matrices were made symmetric and the diagonal was set to zero. To facilitate quantitative comparison across participants, the connectomes were further scaled using inter-subject connection-density normalization following the quantitative streamlines tractography framework (75). Specifically, each participant’s connectome was multiplied by a subject-specific global scaling factor derived from the SIFT2 proportionality coefficient (µ), the median b = 0 intensity within a white-matter mask, the amplitude of the white-matter response function, and voxel volume.

### Functional MRI preprocessing and functional connectivity

The functional images were corrected for slice-timing differences, motion, and signal distortions. Following anatomical T1-coregistration, the functional images were projected to subject-specific cortical surfaces using FreeSurfer v.6.0.1 (73). The time-series were subsequently demeaned and detrended, followed by simultaneous nuisance regression and temporal high-pass filtering (0.009 Hz) so as not to reintroduce nuisance signals (76). Nuisance regression of physiological variables included average white matter, cerebrospinal fluid, and whole-brain time series, six motion parameters, in addition to derivatives, squares, and squared derivatives of each variable (77, 78). To further control for motion, a set of binary spike regressors was included for volumes with framewise displacement > 0.5 mm (79).

Parcelwise functional connectomes were then constructed for each participant and state using Pearson correlations across parcels from the HCP-MMP1 atlas (72), which was projected from fsaverage to individual cortical surfaces using FreeSurfer (73). The resulting correlation coefficients were Fisher z-transformed with matrix diagonals set to zero. Each matrix was adjusted for node volume using the same approach applied to the structural connectomes (74).

To reduce inflation of within-subject FC by shared task coactivation during the movie-watching and working-memory tasks (80, 81), regional leave-one-out group-average time series were computed and regressed out from each participant’s parcelwise time series before connectome creation. This approach targets stimulus-locked signals shared across participants (40, 42, 54) and is conceptually similar to approaches used to separate shared and idiosyncratic components of naturalistic fMRI responses (82, 83). For the working-memory task, connectomes were computed both across the full task run and separately for 1-back, 2-back, and 3-back epochs using condition-specific volume masks.

### CamCAN preprocessing and connectivity estimation

Diffusion preprocessing, tractography, cortical parcellation, and functional image preprocessing followed the same general procedures described above, with the CamCAN-specific steps detailed below. For the DWI data, white-matter fiber orientation distributions were reconstructed using multi-shell multi-tissue constrained spherical deconvolution (MSMT-CSD) (84), implemented in MRtrix3 (59), followed by the same tractography steps described above. For the functional data, nuisance regression combined quadratic detrending, mean white-matter and cerebrospinal-fluid signals, Friston-24 motion regressors, impulse regressors for FD > 0.2 mm and nonsteady-state volumes, and a 0.009-Hz discrete cosine high-pass filter constructed using each run’s TR. Global signal regression and low-pass filtering were not applied. As described above, shared task-related signals were removed using parcelwise leave-one-out group averages before FC estimation. Structural and functional connectivity matrices were then computed following the same methods described above.

### Structure–function coupling

Two complementary approaches were used to estimate SFC. For the whole-brain analyses, SFC was computed separately for each subject and state as the Pearson correlation between log(1 + SC) and FC weights after parcel-volume scaling. Only finite non-zero edges in the upper triangle were included, excluding the diagonal. Regional SFC was estimated for each brain region by correlating its SC profile with its FC profile. Specifically, for each region, the vector of SC weights linking that region to all other regions was correlated with the corresponding vector of FC values in each state. SC weights were log-transformed, and structurally zero, nonfinite, and self-connections were excluded. This yielded one SFC estimate per subject, state, and region, reflecting the degree to which the region’s functional connectivity profile followed its structural connectivity profile.

### Connectivity ratio

As a complementary measure of structure-aligned functional organization, we computed a connectivity ratio (CR) for each participant and state. Whereas SFC estimates the continuous association between SC and FC weights, CR tests whether FC is preferentially expressed along edges with detected structural connections. Structural edges were binarized at SC > 0, defining edges with versus without reconstructed streamline connections. Mean FC was computed separately across structurally connected and structurally undetected edges. CR was subsequently calculated as the relative difference in mean FC between these edge sets, normalized by mean FC among structurally connected edges as shown in Equation (1):

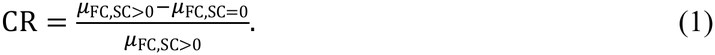

Higher CR values indicate that FC is more strongly concentrated along edges with detected structural connections, whereas lower values indicate weaker preferential alignment of FC and direct structural pathways.

### Functional topology

System segregation was computed from FC using canonical network labels from the Cole-Anticevic partition (85) assigned to HCP-MMP1 parcels (72). For each participant and state, system segregation was calculated the normalized difference between average within-system and between-system connectivity: (within-system FC -between-system FC) / within-system FC. Higher values indicate a more segregated functional configuration.

### Functional stability

Group-level functional stability was estimated from the three group-average FC matrices. For each parcel, Pearson correlations were calculated between its FC profiles for each state pair. Stability was defined as the mean of the three squared correlations minus the standard deviation of their unsquared values, using complete profiles, including their diagonal entries. The stability map was related to mean regional SFC using Spearman correlations and spatial permutation tests. These measures describe similarity across scan conditions rather than temporal stability within a scan.

### Statistical analysis

Analyses were performed in MATLAB R2025b. For both cohorts, the primary state model included state, linear and quadratic age, sex, and mean FD as fixed effects, with a participant-specific random intercept. The mixed models used all eligible state observations, whereas paired differences required both relevant runs. The simplified linear form is shown in Equation (2); the primary model additionally included the quadratic age term:

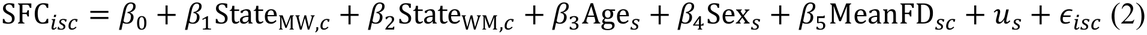

where SFC*isc* denotes the SFC estimate for subject *s*, region or global estimate *i*, and state *c*. StateMW,*c* and StateWM,*c* are dummy variables coding movie-watching and working memory relative to the reference state, resting-state. Age*s*, Sex*s*, and MeanFD*sc* denote chronological age, sex, and state-specific mean framewise displacement, respectively. The term *us* represents the subject-specific random intercept, and *εisc* denotes the residual error.

Models of age-related differences included state interactions with linear and quadratic age to allow for curvature across adulthood. Age associations within each state and differences between state trajectories were evaluated jointly across the two age terms. The simplified interaction form is shown in Equation (3). Working-memory load was coded −1, 0, and 1, with age, age squared, sex, load-specific FD, and a participant intercept included in the model. A state-by-segregation interaction model tested whether the association between segregation and SFC differed across states, and pairwise slope comparisons used two-sided, unadjusted tests.

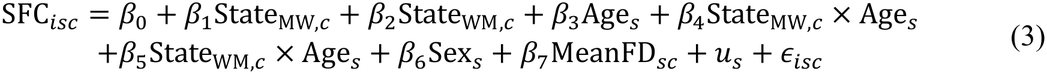

Load was coded −1, 0, and 1, with linear and quadratic age, sex, and within-load standardized FD included as covariates. Segregation and regional state models used standardized FD, retaining the same demographic terms. Regional effects were corrected across 1,080 parcel-by-contrast tests, and network-mean effects across 36 contrasts, using Benjamini–Hochberg corrections. The segregation-adjusted and base models used identical observations and motion covariates. Segregation was standardized across eligible subject-state observations. Reconfiguration was computed as 1 – Pearson correlation between vectorized task and rest FC. Task–rest SFC differences were modeled with the full state-by-reconfiguration-by-age interaction, sex, FD, and participant intercepts. Age was mean centered. Full fixed-effect estimates and additional procedures are provided in the supplementary materials.

The principal resting-state functional gradient was estimated with BrainSpace using cosine similarity, diffusion mapping, and 90% sparsification (86). Spatial correspondence was evaluated using a spin-test with 10,000 rotations (87). Tests were two-sided, with separate FDR correction for the three gradient and stability associations. We additionally resampled participants jointly across states 2,000 times to estimate percentile intervals for differences in spatial correlation.

The age analyses for global SFC used was the original edge-level correlation while network SFC was the participant-level mean of parcelwise correlations within each of the 12 cortical networks. Separate models were fitted to global SFC, every network mean, and each of 360 cortical parcels, using the linear model in Equation (4) and the quadratic model in Equation (5):

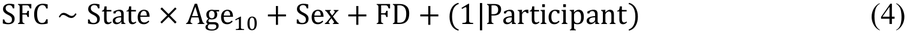

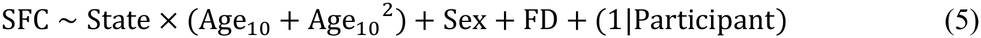

where 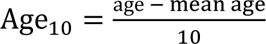. Linear slopes came from the linear-only models. Quadratic coefficients tested curvature, while joint two-degree-of-freedom tests evaluated age association within each state and trajectory differences for every state pair.

## Acknowledgments

Data for the replication analyses were provided by the Cambridge Centre for Ageing and Neuroscience (CamCAN). CamCAN data collection and sharing were funded by the UK Biotechnology and Biological Sciences Research Council (BB/H008217/1), with additional support from the UK Medical Research Council and the University of Cambridge.

## Author contributions

R.P. performed the formal analyses and led manuscript preparation. A.S. supported the analyses and interpretation of the findings and contributed to writing the manuscript. J.P. and R.S.-L. contributed to interpretation of the results, reviewed, and edited the manuscript.

## Data, code, and materials availability

Subject-specific DyNAMiC data are not publicly accessible due to privacy or ethical restrictions but are available upon reasonable request. Access to data by qualified investigators is subject to scientific and ethical review and must comply with the European Union General Data Protection Regulations (GDPR) and all relevant guidelines. The completion of a data transfer agreement signed by an institutional official will be required. Requests should be directed to Alireza Salami. The CamCAN data used in this study were obtained from the Cambridge Centre for Ageing and Neuroscience repository. Access is available through the MRC Cognition and Brain Sciences Unit data portal (https://opendata.mrc-cbu.cam.ac.uk/projects/camcan/), subject to approval of a research proposal and acceptance of the CamCAN data-use agreement. The code used to process the data and perform the analyses reported in this study is available at https://github.com/robinpedersen/SFC_state_differences.

## Supplementary Materials

Supplementary Text

Figs. S1 to S5

Tables S1 to S15

## References

1. C. J. Honey, O. Sporns, L. Cammoun, X. Gigandet, J. P. Thiran, R. Meuli, P. Hagmann, Predicting human resting-state functional connectivity from structural connectivity. Proc. Natl Acad. Sci. USA 106, 2035–2040 (2009). 10.1073/pnas.0811168106

2. H.-J. Park, K. Friston, Structural and functional brain networks: from connections to cognition. Science 342, 1238411 (2013). 10.1126/science.1238411

3. L. E. Suárez, R. D. Markello, R. F. Betzel, B. Mišić, Linking structure and function in macroscale brain networks. Trends Cogn. Sci. 24, 302–315 (2020). 10.1016/j.tics.2020.01.008

4. P. Fotiadis, L. Parkes, K. A. Davis, T. D. Satterthwaite, R. T. Shinohara, D. S. Bassett, Structure-function coupling in macroscale human brain networks. Nat. Rev. Neurosci. 25, 688–704 (2024). 10.1038/s41583-024-00846-6

5. Z.-Q. Liu, B. Vázquez-Rodríguez, R. N. Spreng, B. C. Bernhardt, R. F. Betzel, B. Misic, Time-resolved structure-function coupling in brain networks. Commun. Biol. 5, 532 (2022). 10.1038/s42003-022-03466-x

6. Z. Gu, K. W. Jamison, M. R. Sabuncu, A. Kuceyeski, Heritability and interindividual variability of regional structure-function coupling. Nat. Commun. 12, 4894 (2021). 10.1038/s41467-021-25184-4

7. J. L. Popp, J. A. Thiele, J. Faskowitz, C. Seguin, O. Sporns, K. Hilger, Structural-functional brain network coupling during cognitive demand reveals intelligence-relevant communication strategies. Commun. Biol. 8, 855 (2025). 10.1038/s42003-025-08231-4

8. A. Griffa, E. Amico, R. Liégeois, D. Van De Ville, M. G. Preti, Brain structure-function coupling provides signatures for task decoding and individual fingerprinting. Neuroimage 250, 118970 (2022). 10.1016/j.neuroimage.2022.118970

9. M. W. Cole, J. R. Reynolds, J. D. Power, G. Repovs, A. Anticevic, T. S. Braver, Multi-task connectivity reveals flexible hubs for adaptive task control. Nat. Neurosci. 16, 1348–1355 (2013). 10.1038/nn.3470

10. M. W. Cole, D. S. Bassett, J. D. Power, T. S. Braver, S. E. Petersen, Intrinsic and task-evoked network architectures of the human brain. Neuron 83, 238–251 (2014). 10.1016/j.neuron.2014.05.014

11. J. R. Cohen, M. D’Esposito, The segregation and integration of distinct brain networks and their relationship to cognition. J. Neurosci. 36, 12083–12094 (2016). 10.1523/JNEUROSCI.2965-15.2016

12. F. M. Krienen, B. T. T. Yeo, R. L. Buckner, Reconfigurable task-dependent functional coupling modes cluster around a core functional architecture. Philos. Trans. R. Soc. Lond. B Biol. Sci. 369, 20130526 (2014). 10.1098/rstb.2013.0526

13. J. M. Shine, P. G. Bissett, P. T. Bell, O. Koyejo, J. H. Balsters, K. J. Gorgolewski, C. A. Moodie, R.A. Poldrack, The dynamics of functional brain networks: integrated network states during cognitive task performance. Neuron 92, 544–554 (2016). 10.1016/j.neuron.2016.09.018

14. J. M. Shine, M. Breakspear, P. T. Bell, K. A. Ehgoetz Martens, R. Shine, O. Koyejo, O. Sporns, R. A. Poldrack, Human cognition involves the dynamic integration of neural activity and neuromodulatory systems. Nat. Neurosci. 22, 289–296 (2019). 10.1038/s41593-018-0312-0

15. G. S. Wig, Segregated systems of human brain networks. Trends Cogn. Sci. 21, 981–996 (2017). 10.1016/j.tics.2017.09.006

16. A. Avena-Koenigsberger, B. Mišić, O. Sporns, Communication dynamics in complex brain networks. Nat. Rev. Neurosci. 19, 17–33 (2018). 10.1038/nrn.2017.149

17. J. Goñi, M. P. van den Heuvel, A. Avena-Koenigsberger, N. Velez de Mendizabal, R. F. Betzel, A. Griffa, P. Hagmann, B. Corominas-Murtra, J.-P. Thiran, O. Sporns, Resting-brain functional connectivity predicted by analytic measures of network communication. Proc. Natl Acad. Sci. USA 111, 833–838 (2014). 10.1073/pnas.1315529111

18. C. Seguin, O. Sporns, A. Zalesky, Brain network communication: concepts, models and applications. Nat. Rev. Neurosci. 24, 557–574 (2023). 10.1038/s41583-023-00718-5

19. F. Zamani Esfahlani, J. Faskowitz, J. Slack, B. Mišić, R. F. Betzel, Local structure-function relationships in human brain networks across the lifespan. Nat. Commun. 13, 2053 (2022). 10.1038/s41467-022-29770-y

20. G. L. Baum, Z. Cui, D. R. Roalf, R. Ciric, R. F. Betzel, B. Larsen, M. Cieslak, P. A. Cook, C. H. Xia, T. M. Moore, K. Ruparel, D. J. Oathes, A. F. Alexander-Bloch, R. T. Shinohara, A. Raznahan, R. E. Gur, R. C. Gur, D. S. Bassett, T. D. Satterthwaite, Development of structure-function coupling in human brain networks during youth. Proc. Natl Acad. Sci. USA 117, 771–778 (2020). 10.1073/pnas.1912034117

21. M. G. Preti, D. Van De Ville, Decoupling of brain function from structure reveals regional behavioral specialization in humans. Nat. Commun. 10, 4747 (2019). 10.1038/s41467-019-12765-7

22. B. Vázquez-Rodríguez, L. E. Suárez, R. D. Markello, G. Shafiei, C. Paquola, P. Hagmann, M. P. van den Heuvel, B. C. Bernhardt, R. N. Spreng, B. Misic, Gradients of structure-function tethering across neocortex. Proc. Natl Acad. Sci. USA 116, 21219–21227 (2019). 10.1073/pnas.1903403116

23. R. L. Buckner, F. M. Krienen, The evolution of distributed association networks in the human brain.Trends Cogn. Sci. 17, 648–665 (2013). 10.1016/j.tics.2013.09.017

24. M.-M. Mesulam, Large-scale neurocognitive networks and distributed processing for attention, language, and memory. Ann. Neurol. 28, 597–613 (1990). 10.1002/ana.410280502

25. M. P. van den Heuvel, R. C. W. Mandl, R. S. Kahn, H. E. Hulshoff Pol, Functionally linked resting-state networks reflect the underlying structural connectivity architecture of the human brain. Hum. Brain Mapp. 30, 3127–3141 (2009). 10.1002/hbm.20737

26. P. B. Kemmer, Y. Wang, F. D. Bowman, H. Mayberg, Y. Guo, Evaluating the strength of structural connectivity underlying brain functional networks. Brain Connect. 8, 579–594 (2018). 10.1089/brain.2018.0615

27. M. Fukushima, R. F. Betzel, Y. He, M. P. van den Heuvel, X.-N. Zuo, O. Sporns, Structure–function relationships during segregated and integrated network states of human brain functional connectivity. Brain Struct. Funct. 223, 1091–1106 (2018). 10.1007/s00429-017-1539-3

28. Y. Osmanlıoğlu, B. Tunç, D. Parker, M. A. Elliott, G. L. Baum, R. Ciric, T. D. Satterthwaite, R. E. Gur, R. C. Gur, R. Verma, System-level matching of structural and functional connectomes in the human brain. Neuroimage 199, 93–104 (2019). 10.1016/j.neuroimage.2019.05.064

29. B. Mišić, R. F. Betzel, M. A. de Reus, M. P. van den Heuvel, M. G. Berman, A. R. McIntosh, O. Sporns, Network-level structure-function relationships in human neocortex. Cereb. Cortex 26, 3285–3296 (2016). 10.1093/cercor/bhw089

30. J. Zimmermann, P. Ritter, K. Shen, S. Rothmeier, M. Schirner, A. R. McIntosh, Structural architecture supports functional organization in the human aging brain at a regionwise and network level. Hum. Brain Mapp. 37, 2645–2661 (2016). 10.1002/hbm.23200

31. A. M. Hermundstad, D. S. Bassett, K. S. Brown, E. M. Aminoff, D. Clewett, S. Freeman, A. Frithsen, A. Johnson, C. M. Tipper, M. B. Miller, S. T. Grafton, J. M. Carlson, Structural foundations of resting-state and task-based functional connectivity in the human brain. Proc. Natl Acad. Sci. USA 110, 6169–6174 (2013). 10.1073/pnas.1219562110

32. A. M. Hermundstad, K. S. Brown, D. S. Bassett, E. M. Aminoff, A. Frithsen, A. Johnson, C. M. Tipper, M. B. Miller, S. T. Grafton, J. M. Carlson, Structurally-constrained relationships between cognitive states in the human brain. PLoS Comput. Biol. 10, e1003591 (2014). 10.1371/journal.pcbi.1003591

33. K. Nordin, T. Gorbach, R. Pedersen, V. Panes Lundmark, J. Johansson, M. Andersson, C. McNulty, K. Riklund, A. Wåhlin, G. Papenberg, G. Kalpouzos, L. Bäckman, A. Salami, DyNAMiC: a prospective longitudinal study of dopamine and brain connectomes: a new window into cognitive aging. J. Neurosci. Res. 100, 1296–1320 (2022). 10.1002/jnr.25039

34. M. A. Shafto, L. K. Tyler, M. Dixon, J. R. Taylor, J. B. Rowe, R. Cusack, A. J. Calder, W. D. Marslen-Wilson, J. Duncan, T. Dalgleish, R. N. Henson, C. Brayne, F. E. Matthews, Cam-CAN, The Cambridge Centre for Ageing and Neuroscience (Cam-CAN) study protocol: a cross-sectional, lifespan, multidisciplinary examination of healthy cognitive ageing. BMC Neurol. 14, 204 (2014). 10.1186/s12883-014-0204-1

35. J. R. Taylor, N. Williams, R. Cusack, T. Auer, M. A. Shafto, M. Dixon, L. K. Tyler, Cam-CAN, R. N. Henson, The Cambridge Centre for Ageing and Neuroscience (Cam-CAN) data repository: structural and functional MRI, MEG, and cognitive data from a cross-sectional adult lifespan sample. Neuroimage 144, 262–269 (2017). 10.1016/j.neuroimage.2015.09.018

36. M. D. Fox, M. E. Raichle, Spontaneous fluctuations in brain activity observed with functional magnetic resonance imaging. Nat. Rev. Neurosci. 8, 700–711 (2007). 10.1038/nrn2201

37. A. M. Morcom, P. C. Fletcher, Does the brain have a baseline? Why we should be resisting a rest. Neuroimage 37, 1073–1082 (2007). 10.1016/j.neuroimage.2006.09.013

38. M. D. Greicius, K. Supekar, V. Menon, R. F. Dougherty, Resting-state functional connectivity reflects structural connectivity in the default mode network. Cereb. Cortex 19, 72–78 (2009). 10.1093/cercor/bhn059

39. J. Gonzalez-Castillo, J. W. Y. Kam, C. W. Hoy, P. A. Bandettini, How to interpret resting-state fMRI: ask your participants. J. Neurosci. 41, 1130–1141 (2021). 10.1523/JNEUROSCI.1786-20.2020

40. U. Hasson, Y. Nir, I. Levy, G. Fuhrmann, R. Malach, Intersubject synchronization of cortical activity during natural vision. Science 303, 1634–1640 (2004). 10.1126/science.1089506

41. V. Betti, S. Della Penna, F. de Pasquale, D. Mantini, L. Marzetti, G. L. Romani, M. Corbetta, Natural scenes viewing alters the dynamics of functional connectivity in the human brain. Neuron 79, 782–797 (2013). 10.1016/j.neuron.2013.06.022

42. E. Simony, C. J. Honey, J. Chen, O. Lositsky, Y. Yeshurun, A. Wiesel, U. Hasson, Dynamic reconfiguration of the default mode network during narrative comprehension. Nat. Commun. 7, 12141 (2016). 10.1038/ncomms12141

43. T. Vanderwal, J. Eilbott, E. S. Finn, R. C. Craddock, A. Turnbull, F. X. Castellanos, Individual differences in functional connectivity during naturalistic viewing conditions. Neuroimage 157, 521–530 (2017). 10.1016/j.neuroimage.2017.06.027

44. S. A. Nastase, A. Goldstein, U. Hasson, Keep it real: rethinking the primacy of experimental control in cognitive neuroscience. Neuroimage 222, 117254 (2020). 10.1016/j.neuroimage.2020.117254

45. S. Sonkusare, M. Breakspear, C. Guo, Naturalistic stimuli in neuroscience: critically acclaimed. Trends Cogn. Sci. 23, 699–714 (2019). 10.1016/j.tics.2019.05.004

46. P. Fotiadis, A. F. T. Arnsten, L. Parkes, T. D. Satterthwaite, R. T. Shinohara, D. S. Bassett, Biological substrates of structure-function coupling in brain networks. Neurosci. Biobehav. Rev. 184, 106581 (2026). 10.1016/j.neubiorev.2026.106581

47. V. Subramani, G. Lioi, K. Jerbi, N. Farrugia, Structure-function coupling and decoupling during movie watching and resting state: novel insights bridging EEG and structural imaging. Imaging Neurosci. 3, imag_a_00448 (2025). 10.1162/imag_a_00448

48. R. F. Betzel, L. Byrge, Y. He, J. Goñi, X.-N. Zuo, O. Sporns, Changes in structural and functional connectivity among resting-state networks across the human lifespan. Neuroimage 102, 345–357 (2014). 10.1016/j.neuroimage.2014.07.067

49. R. Pedersen, L. Geerligs, M. Andersson, T. Gorbach, B. Avelar-Pereira, A. Wåhlin, A. Rieckmann, L. Nyberg, A. Salami, When functional blurring becomes deleterious: reduced system segregation is associated with less white matter integrity and cognitive decline in aging. Neuroimage 242, 118449 (2021). 10.1016/j.neuroimage.2021.118449

50. X. Liu, L. K. Tyler, Cam-CAN, S. W. Davis, J. B. Rowe, K. A. Tsvetanov, Cognition’s dependence on functional network integrity with age is conditional on structural network integrity. Neurobiol. Aging 129, 195–208 (2023). 10.1016/j.neurobiolaging.2023.06.001

51. C. Guichet, S. Achard, M. Mermillod, M. Baciu, Structure-function dynamics in healthy cognitive aging: a graph signal processing approach. Brain Struct. Funct. 231, 85 (2026). 10.1007/s00429-026-03130-8

52. D. K. Jones, T. R. Knösche, R. Turner, White matter integrity, fiber count, and other fallacies: the do’s and don’ts of diffusion MRI. Neuroimage 73, 239–254 (2013). 10.1016/j.neuroimage.2012.06.081

53. K. H. Maier-Hein, P. F. Neher, J.-C. Houde, M.-A. Côté, E. Garyfallidis, J. Zhong, M. Chamberland, F.-C. Yeh, Y.-C. Lin, Q. Ji, W. E. Reddick, J. O. Glass, D. Q. Chen, Y. Feng, C. Gao, Y. Wu, J. Ma, R. He, Q. Li, C.-F. Westin, S. Deslauriers-Gauthier, J. O. O. González, M. Paquette, S. St-Jean, G. Girard, F. Rheault, J. Sidhu, C. M. W. Tax, F. Guo, H. Y. Mesri, S. Dávid, M. Froeling, A. M. Heemskerk, A. Leemans, A. Boré, B. Pinsard, C. Bedetti, M. Desrosiers, S. Brambati, J. Doyon, A. Sarica, R. Vasta, A. Cerasa, A. Quattrone, J. Yeatman, A. R. Khan, W. Hodges, S. Alexander, D. Romascano, M. Barakovic, A. Auría, O. Esteban, A. Lemkaddem, J.-P. Thiran, H. E. Cetingul, B. L. Odry, B. Mailhe, M. S. Nadar, F. Pizzagalli, G. Prasad, J. E. Villalon-Reina, J. Galvis, P. M. Thompson, F. D. S. Requejo, P. L. Laguna, L. M. Lacerda, R. Barrett, F. Dell’Acqua, M. Catani, L. Petit, E. Caruyer, A. Daducci, T. B. Dyrby, T. Holland-Letz, C. C. Hilgetag, B. Stieltjes, M. Descoteaux, The challenge of mapping the human connectome based on diffusion tractography. Nat. Commun. 8, 1349 (2017). 10.1038/s41467-017-01285-x

54. S. A. Nastase, V. Gazzola, U. Hasson, C. Keysers, Measuring shared responses across subjects using intersubject correlation. Soc. Cogn. Affect. Neurosci. 14, 667–685 (2019). 10.1093/scan/nsz037

55. E. S. Finn, P. A. Bandettini, Movie-watching outperforms rest for functional connectivity-based prediction of behavior. Neuroimage 235, 117963 (2021). 10.1016/j.neuroimage.2021.117963

56. R. M. Hutchison, T. Womelsdorf, E. A. Allen, P. A. Bandettini, V. D. Calhoun, M. Corbetta, S. Della Penna, J. H. Duyn, G. H. Glover, J. Gonzalez-Castillo, D. A. Handwerker, S. Keilholz, V. Kiviniemi, D. A. Leopold, F. de Pasquale, O. Sporns, M. Walter, C. Chang, Dynamic functional connectivity: promise, issues, and interpretations. Neuroimage 80, 360–378 (2013). 10.1016/j.neuroimage.2013.05.079

57. M. G. Preti, T. A. W. Bolton, D. Van De Ville, The dynamic functional connectome: state-of-the-art and perspectives. Neuroimage 160, 41–54 (2017). 10.1016/j.neuroimage.2016.12.061

58. Y. Yang, Z. Zheng, L. Liu, H. Zheng, Y. Zhen, Y. Zheng, X. Wang, S. Tang, Enhanced brain structure-function tethering in transmodal cortex revealed by high-frequency eigenmodes. Nat. Commun. 14, 6744 (2023). 10.1038/s41467-023-42053-4

59. J.-D. Tournier, R. Smith, D. Raffelt, R. Tabbara, T. Dhollander, M. Pietsch, D. Christiaens, B. Jeurissen, C.-H. Yeh, A. Connelly, MRtrix3: a fast, flexible and open software framework for medical image processing and visualisation. Neuroimage 202, 116137 (2019). 10.1016/j.neuroimage.2019.116137

60. J. Veraart, E. Fieremans, D. S. Novikov, Diffusion MRI noise mapping using random matrix theory. Magn. Reson. Med. 76, 1582–1593 (2016). 10.1002/mrm.26059

61. J. Veraart, D. S. Novikov, D. Christiaens, B. Ades-aron, J. Sijbers, E. Fieremans, Denoising of diffusion MRI using random matrix theory. Neuroimage 142, 394–406 (2016). 10.1016/j.neuroimage.2016.08.016

62. E. Kellner, B. Dhital, V. G. Kiselev, M. Reisert, Gibbs-ringing artifact removal based on local subvoxel-shifts. Magn. Reson. Med. 76, 1574–1581 (2016). 10.1002/mrm.26054

63. J. L. R. Andersson, S. Skare, J. Ashburner, How to correct susceptibility distortions in spin-echo echo-planar images: application to diffusion tensor imaging. Neuroimage 20, 870–888 (2003). 10.1016/S1053-8119(03)00336-7

64. J. L. R. Andersson, S. N. Sotiropoulos, An integrated approach to correction for off-resonance effects and subject movement in diffusion MR imaging. Neuroimage 125, 1063–1078 (2016). 10.1016/j.neuroimage.2015.10.019

65. N. J. Tustison, B. B. Avants, P. A. Cook, Y. Zheng, A. Egan, P. A. Yushkevich, J. C. Gee, N4ITK: improved N3 bias correction. IEEE Trans. Med. Imaging 29, 1310–1320 (2010). 10.1109/TMI.2010.2046908

66. T. Dhollander, D. Raffelt, A. Connelly, Unsupervised 3-tissue response function estimation from single-shell or multi-shell diffusion MR data without a co-registered T1 image. In ISMRM Workshop on Breaking the Barriers of Diffusion MRI 5 (2016).

67. T. Dhollander, R. Mito, D. Raffelt, A. Connelly, Improved white matter response function estimation for 3-tissue constrained spherical deconvolution. In Proceedings of the 27th Annual Meeting of the International Society for Magnetic Resonance in Medicine 555 (2019).

68. T. Dhollander, A. Connelly, A novel iterative approach to reap the benefits of multi-tissue CSD from just single-shell (+b=0) diffusion MRI data. In Proceedings of the 24th Annual Meeting of the International Society for Magnetic Resonance in Medicine 3010 (2016).

69. R. E. Smith, J.-D. Tournier, F. Calamante, A. Connelly, Anatomically-constrained tractography: improved diffusion MRI streamlines tractography through effective use of anatomical information. Neuroimage 62, 1924–1938 (2012). 10.1016/j.neuroimage.2012.06.005

70. J.-D. Tournier, F. Calamante, A. Connelly, Improved probabilistic streamlines tractography by 2nd order integration over fibre orientation distributions. In Proceedings of the 18th Annual Meeting of the International Society for Magnetic Resonance in Medicine 1670 (2010).

71. R. E. Smith, J.-D. Tournier, F. Calamante, A. Connelly, SIFT2: enabling dense quantitative assessment of brain white matter connectivity using streamlines tractography. Neuroimage 119, 338–351 (2015). 10.1016/j.neuroimage.2015.06.092

72. M. F. Glasser, T. S. Coalson, E. C. Robinson, C. D. Hacker, J. Harwell, E. Yacoub, K. Ugurbil, J. Andersson, C. F. Beckmann, M. Jenkinson, S. M. Smith, D. C. Van Essen, A multi-modal parcellation of human cerebral cortex. Nature 536, 171–178 (2016). 10.1038/nature18933

73. B. Fischl, FreeSurfer. Neuroimage 62, 774–781 (2012). 10.1016/j.neuroimage.2012.01.021

74. P. Hagmann, L. Cammoun, X. Gigandet, R. Meuli, C. J. Honey, V. J. Wedeen, O. Sporns, Mapping the structural core of human cerebral cortex. PLoS Biol. 6, e159 (2008). 10.1371/journal.pbio.0060159

75. R. E. Smith, D. Raffelt, J.-D. Tournier, A. Connelly, Quantitative streamlines tractography: methods and inter-subject normalisation. Apert. Neuro 2, 1–25 (2022). 10.52294/ApertureNeuro.2022.2.NEOD9565

76. M. N. Hallquist, K. Hwang, B. Luna, The nuisance of nuisance regression: spectral misspecification in a common approach to resting-state fMRI preprocessing reintroduces noise and obscures functional connectivity. Neuroimage 82, 208–225 (2013). 10.1016/j.neuroimage.2013.05.116

77. T. D. Satterthwaite, D. H. Wolf, J. Loughead, K. Ruparel, M. A. Elliott, H. Hakonarson, R. C. Gur, R. E. Gur, Impact of in-scanner head motion on multiple measures of functional connectivity: relevance for studies of neurodevelopment in youth. Neuroimage 60, 623–632 (2012). 10.1016/j.neuroimage.2011.12.063

78. R. Ciric, D. H. Wolf, J. D. Power, D. R. Roalf, G. L. Baum, K. Ruparel, R. T. Shinohara, M. A. Elliott, S. B. Eickhoff, C. Davatzikos, R. C. Gur, R. E. Gur, D. S. Bassett, T. D. Satterthwaite, Benchmarking of participant-level confound regression strategies for the control of motion artifact in studies of functional connectivity. Neuroimage 154, 174–187 (2017). 10.1016/j.neuroimage.2017.03.020

79. J. D. Power, K. A. Barnes, A. Z. Snyder, B. L. Schlaggar, S. E. Petersen, Spurious but systematic correlations in functional connectivity MRI networks arise from subject motion. Neuroimage 59, 2142–2154 (2012). 10.1016/j.neuroimage.2011.10.018

80. M. W. Cole, T. Ito, D. Schultz, R. Mill, R. Chen, C. Cocuzza, Task activations produce spurious but systematic inflation of task functional connectivity estimates. Neuroimage 189, 1–18 (2019). 10.1016/j.neuroimage.2018.12.054

81. R. Masharipov, I. Knyazeva, A. Korotkov, D. Cherednichenko, M. Kireev, Comparison of whole-brain task-modulated functional connectivity methods for fMRI task connectomics. Commun. Biol. 7, 1402 (2024). 10.1038/s42003-024-07088-3

82. C. J. Wild, A. C. Linke, L. Zubiaurre-Elorza, C. Herzmann, H. Duffy, V. K. Han, D. S. C. Lee, R. Cusack, Adult-like processing of naturalistic sounds in auditory cortex by 3- and 9-month old infants. Neuroimage 157, 623–634 (2017). 10.1016/j.neuroimage.2017.06.038

83. R. Tansey, K. Graff, C. S. Rohr, D. Dimond, A. Ip, S. Yin, D. Dewey, S. Bray, Functional MRI responses to naturalistic stimuli are increasingly typical across early childhood. Dev. Cogn. Neurosci. 62, 101268 (2023). 10.1016/j.dcn.2023.101268

84. B. Jeurissen, J.-D. Tournier, T. Dhollander, A. Connelly, J. Sijbers, Multi-tissue constrained spherical deconvolution for improved analysis of multi-shell diffusion MRI data. Neuroimage 103, 411–426 (2014). 10.1016/j.neuroimage.2014.07.061

85. J. L. Ji, M. Spronk, K. Kulkarni, G. Repovš, A. Anticevic, M. W. Cole, Mapping the human brain’s cortical-subcortical functional network organization. Neuroimage 185, 35–57 (2019). 10.1016/j.neuroimage.2018.10.006

86. R. Vos de Wael, O. Benkarim, C. Paquola, S. Lariviere, J. Royer, S. Tavakol, T. Xu, S.-J. Hong, G. Langs, S. Valk, B. Misic, M. Milham, D. Margulies, J. Smallwood, B. C. Bernhardt, BrainSpace: a toolbox for the analysis of macroscale gradients in neuroimaging and connectomics datasets. Commun. Biol. 3, 103 (2020). 10.1038/s42003-020-0794-7

87. A. F. Alexander-Bloch, H. Shou, S. Liu, T. D. Satterthwaite, D. C. Glahn, R. T. Shinohara, S. N. Vandekar, A. Raznahan, On testing for spatial correspondence between maps of human brain structure and function. Neuroimage 178, 540–551 (2018). 10.1016/j.neuroimage.2018.05.070

