## Supplementary Materials for "State-dependent structure-function coupling reflects dynamic engagement of the brain’s structural scaffold"

Robin Pedersen et al.

**This PDF file includes**

Supplementary Text

Figs. S1 to S5

Tables S1 to S15

**Supplementary Text**

**State differences in structural-functional coupling in DyNAMiC and CamCAN**

The following tables report the models supporting the main analyses. DyNAMiC contributes 525 observations from 177 participants, and CamCAN contributes 1153 observations from 546 participants. Models use all eligible runs: Dynamic: resting-state *n* = 176; movie-watching, *n* = 174, and working memory, *n* = 175). CamCAN: resting-state, *n* = 271; movie-watching, *n* = 536, and sensorimotor task, *n* = 346. Tables report fixed effects, two-sided coefficient tests, and 95% confidence intervals. Participant-specific random intercepts account for repeated observations. The reference categories and covariates are specified below each table.

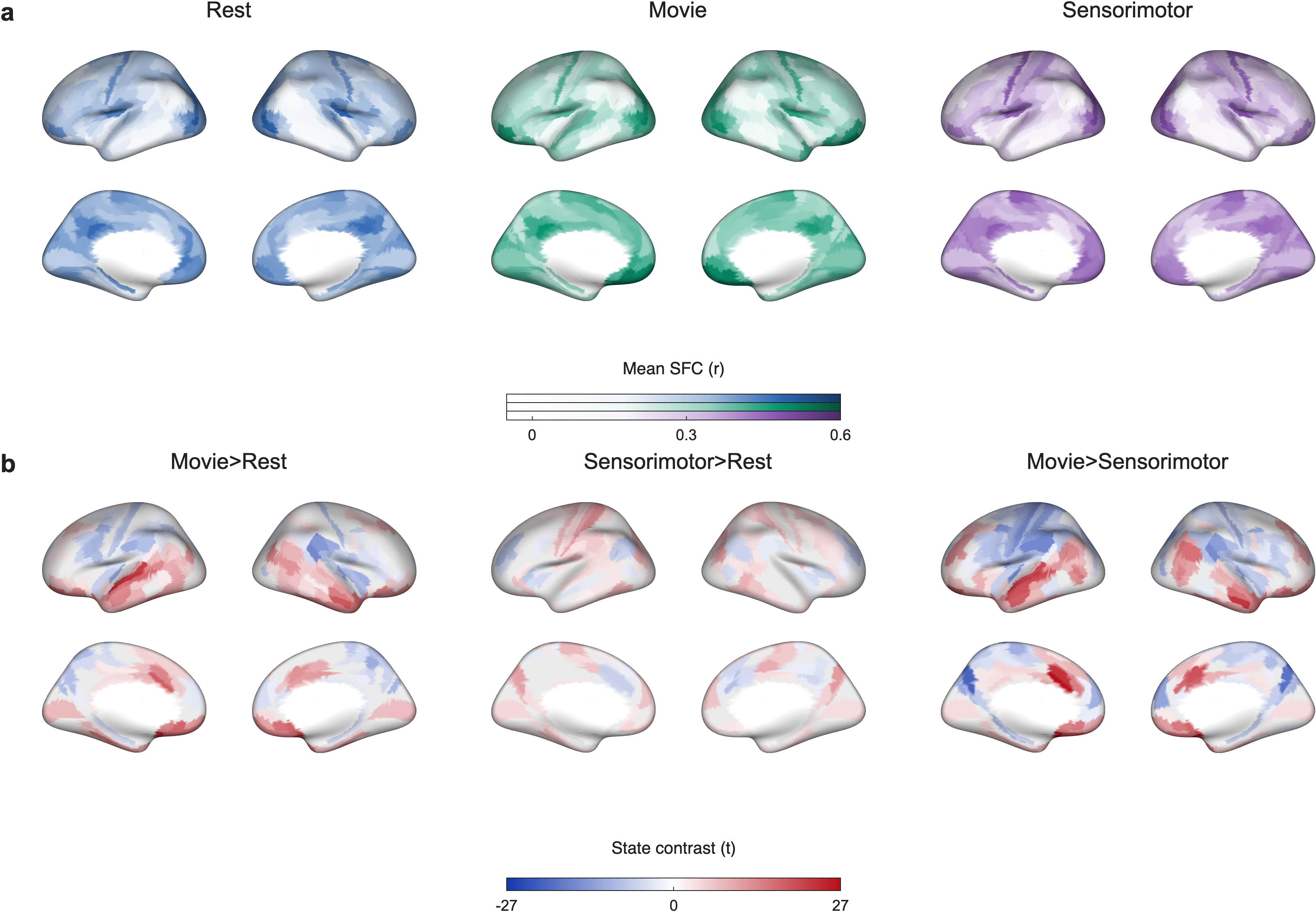

**Fig. S1. Regional structure–function coupling in CamCAN.**

(a) Mean regional SFC during rest, movie-watching, and the sensorimotor task, shown on a common scale. (b) Corresponding state contrasts, with blue–white–red *t*-statistics; positive values indicate higher SFC in the first named state. Only parcels surviving FDR correction across all 1,080 regional comparisons are colored; other parcels are gray. Models include state, age, sex, and mean FD, with a participant-specific intercept.

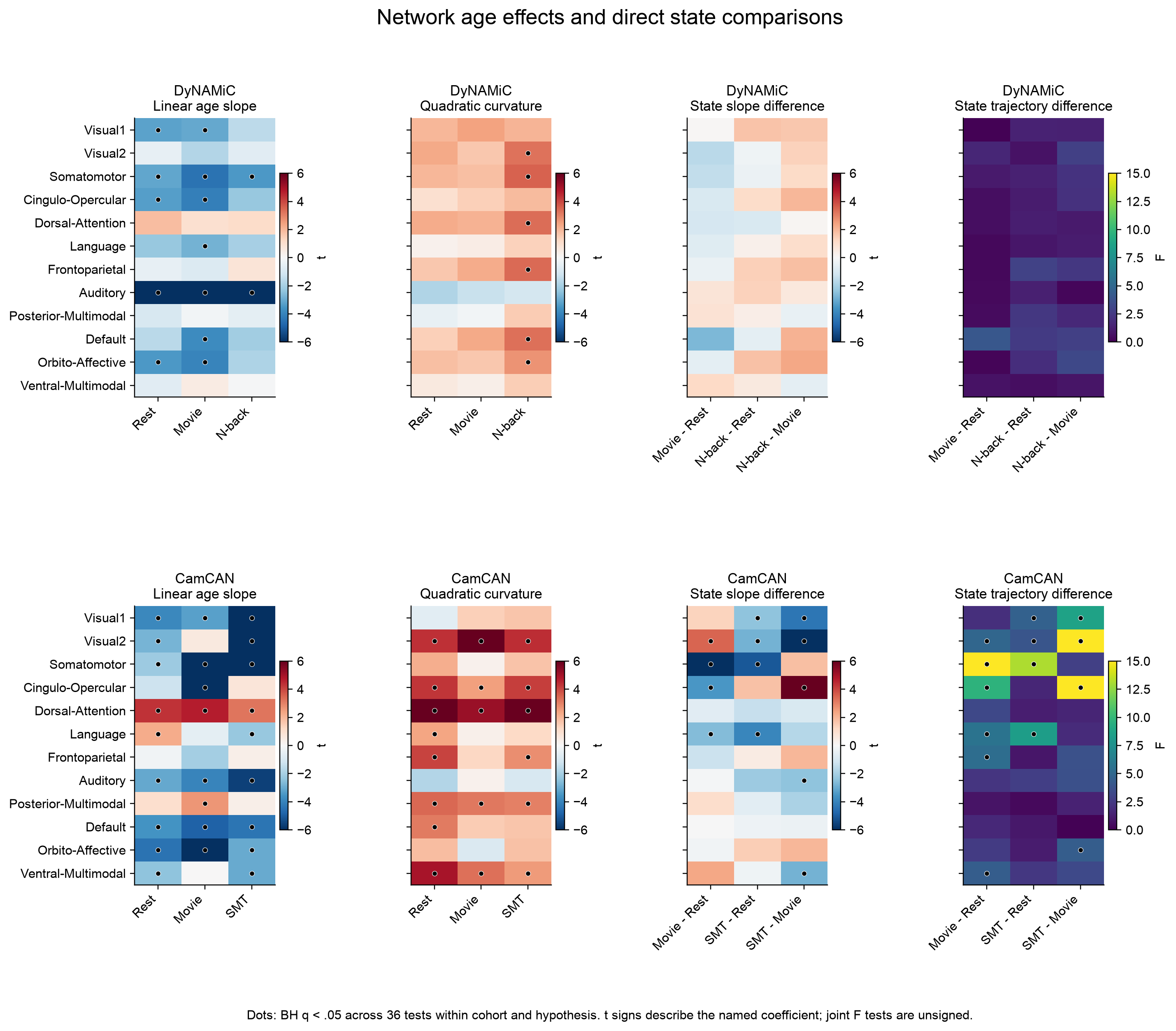

**Fig. S2. Network age associations and state comparisons.**

Network-level age associations and state comparisons in DyNAMiC (top row) and CamCAN (bottom row), shown for all 12 networks. The first three columns show signed *t*-statistics for linear age slopes, quadratic curvature, and between-state differences in linear age slopes; the fourth shows unsigned *F*-statistics for joint differences in linear and quadratic age terms between states. Dots indicate FDR-adjusted *p*-values < .05 across 36 tests within each cohort. Positive curvature indicates convexity rather than a uniformly positive age slope. SMT denotes the sensorimotor task.

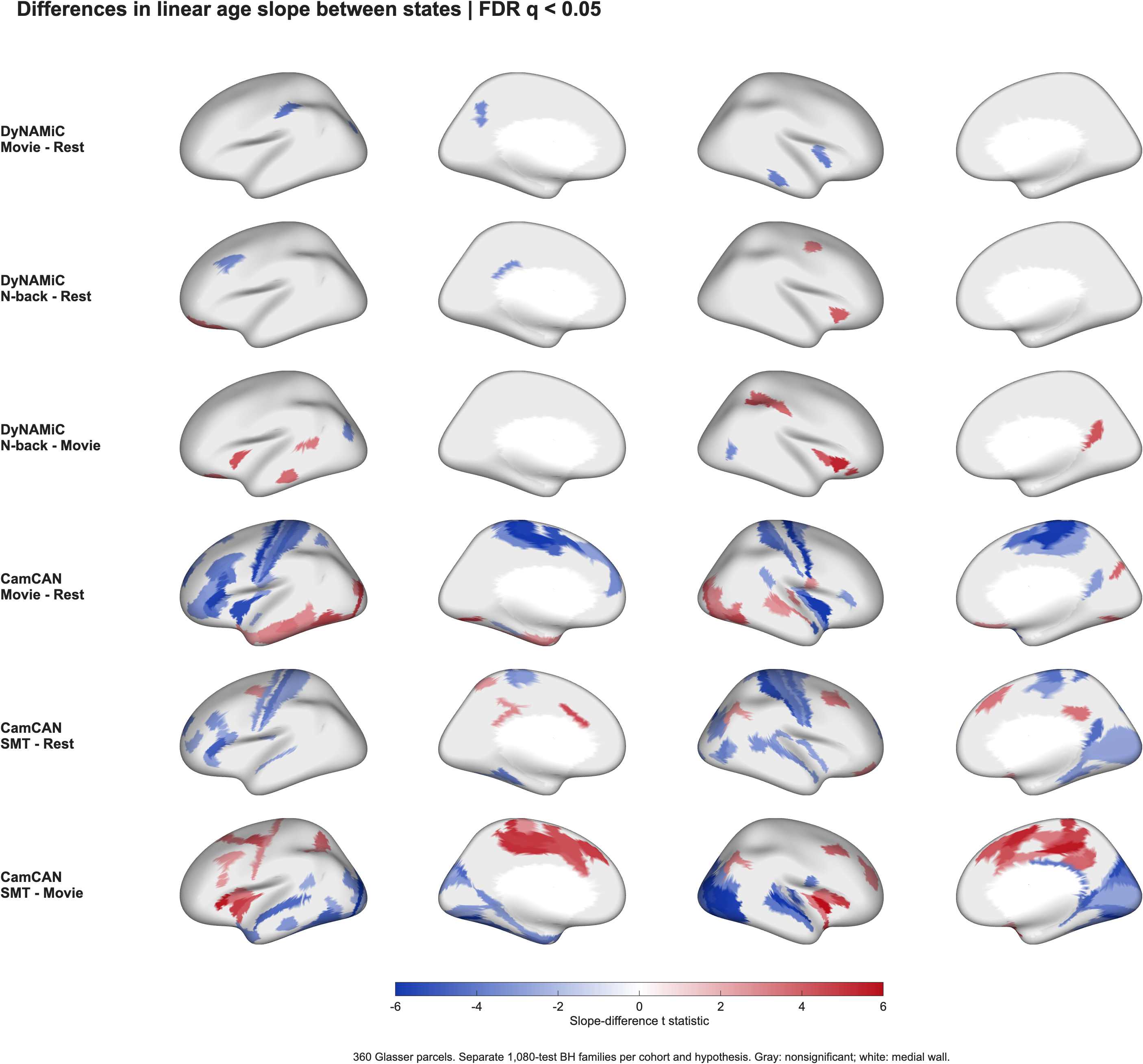

**Fig. S3. Regional differences in linear age slopes.**

State differences in regional linear age associations in DyNAMiC and CamCAN, shown as blue–white–red *t*-statistics from mixed models. Positive values indicate a more positive age slope in the first named state. Only parcels surviving FDR correction across all 1,080 regional comparisons (360 parcels × 3 state pairs) within each cohort are colored; other parcels are gray. SMT denotes the sensorimotor task.

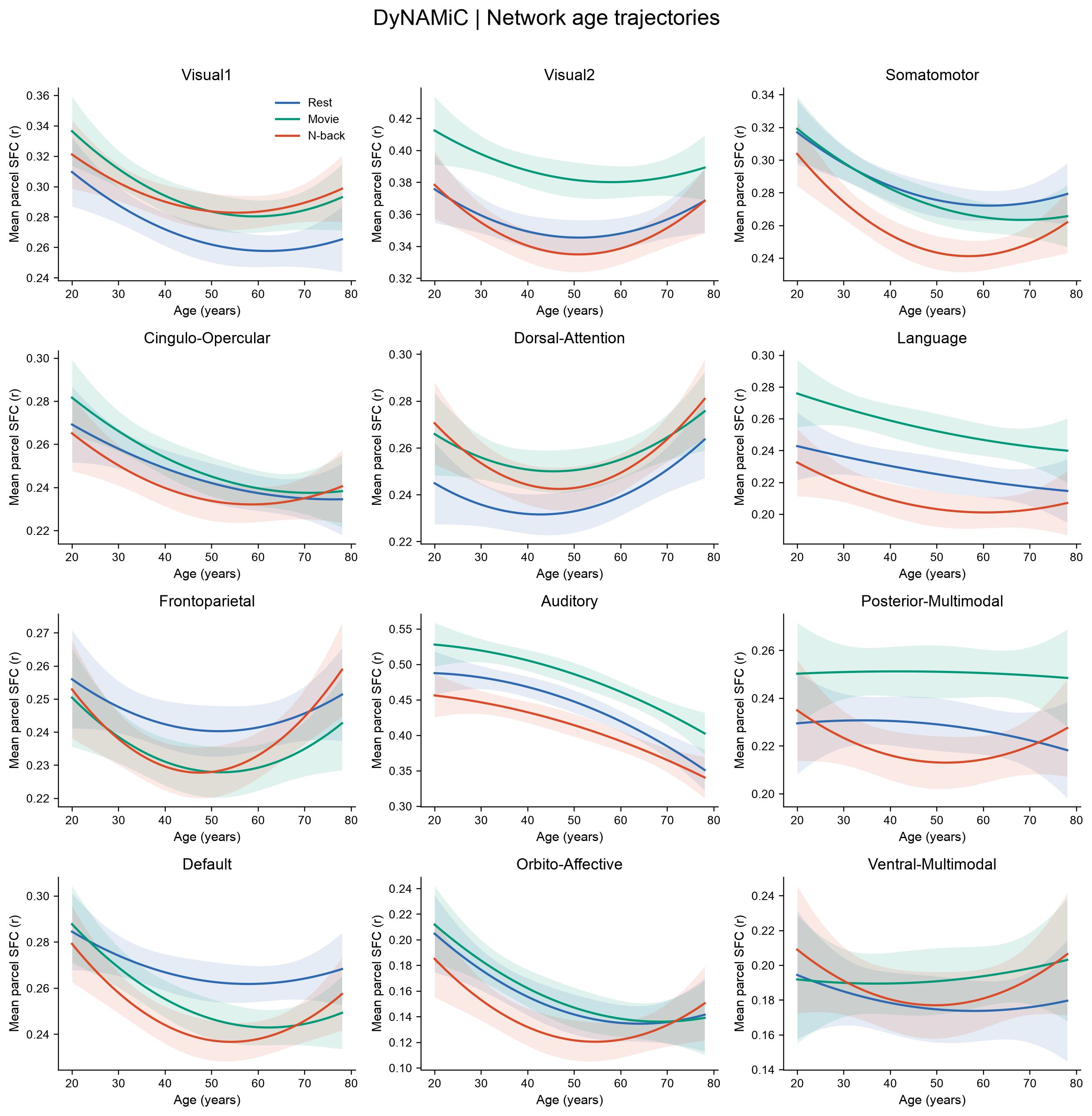

**Fig. S4. Network-level age trajectories in DyNAMiC.**

Age trajectories of mean SFC in all 12 networks during rest, movie-watching, and working memory. Lines show quadratic mixed-model predictions, adjusted to pooled mean motion and observed sex proportions; shading shows pointwise 95% confidence intervals. Curves span the observed age range and are displayed irrespective of significance. Separate linear and quadratic age tests are reported in Table S14.

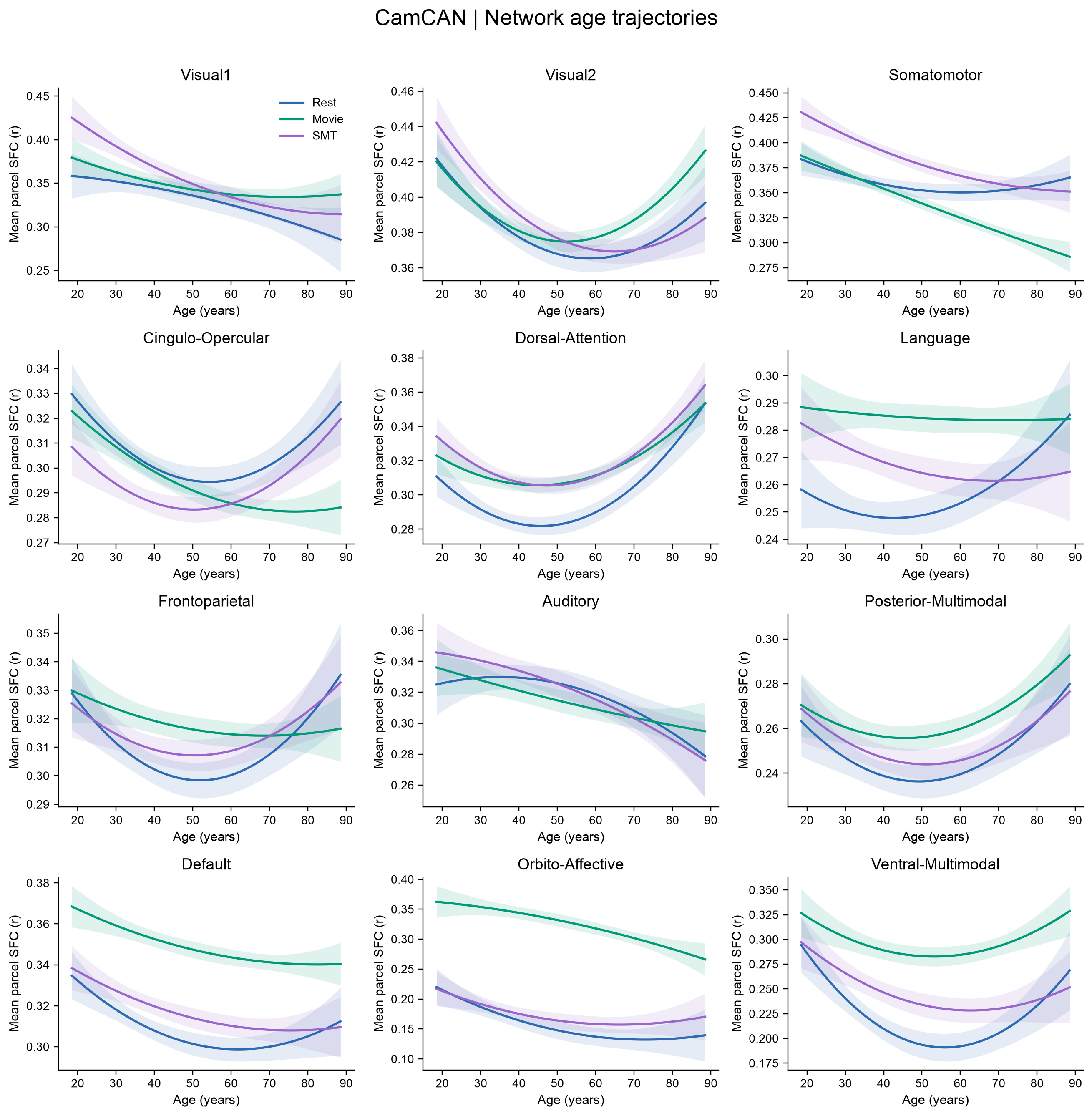

**Fig. S5. Network-level age trajectories in CamCAN.**

Age trajectories of mean SFC in all 12 networks during rest, movie-watching, and the sensorimotor task. Lines show quadratic mixed-model predictions, adjusted for motion using the CamCAN model specification and observed sex proportions; shading shows pointwise 95% confidence intervals. Curves span the observed age range and are displayed irrespective of significance. Separate linear and quadratic age tests are reported in Table S15.

**Table S1. DyNAMiC primary state model**

| **Predictor** | **Estimate** | **SE** | **95% CI** | ***t*** | ***df*** | ***p*** |
| --- | --- | --- | --- | --- | --- | --- |
| Intercept | 0.2635 | 0.0049 | [0.2539, 0.2731] | 53.98 | 518 | <.001 |
| Movie vs. rest | 0.0090 | 0.0022 | [0.0046, 0.0134] | 4.04 | 518 | <.001 |
| N-back vs. rest | −0.0127 | 0.0022 | [−0.0171, −0.0084] | −5.72 | 518 | <.001 |
| Age | −0.0068 | 0.0024 | [−0.0115, −0.0021] | −2.84 | 518 | .005 |
| Sex (female vs. male) | 0.0075 | 0.0046 | [−0.0015, 0.0166] | 1.63 | 518 | .104 |
| Mean FD | −0.0169 | 0.0016 | [−0.0200, −0.0137] | −10.41 | 518 | <.001 |
| Age² | 0.0064 | 0.0027 | [0.0011, 0.0116] | 2.37 | 518 | .018 |

Note: DyNAMiC: 525 eligible observations from 177 participants. Linear mixed model with participant-specific intercepts; rest and male sex are the reference categories. Age is mean centered within the cohort and divided by the cohort SD, with Age² its square. Omnibus state test: *F*(2,518) = 47.89, *p* < 0.001. *p*-values are two-sided and unadjusted.

**Table S2. DyNAMiC connectivity ratio**

| **Predictor** | **Estimate** | **SE** | **95% CI** | ***t*** | ***df*** | ***p*** |
| --- | --- | --- | --- | --- | --- | --- |
| Intercept | 0.1274 | 0.0155 | [0.0970, 0.1578] | 8.23 | 518 | <.001 |
| Movie vs. rest | 0.0111 | 0.0094 | [−0.0074, 0.0297] | 1.18 | 518 | .239 |
| N-back vs. rest | −0.0283 | 0.0094 | [−0.0467, −0.0098] | −3.01 | 518 | .003 |
| Age | 0.0312 | 0.0084 | [0.0146, 0.0478] | 3.70 | 518 | <.001 |
| Sex (female vs. male) | 0.0284 | 0.0161 | [−0.0032, 0.0601] | 1.77 | 518 | .078 |
| Mean FD | 0.0111 | 0.0064 | [−0.0015, 0.0236] | 1.73 | 518 | .084 |
| Age² | 0.0316 | 0.0094 | [0.0132, 0.0500] | 3.37 | 518 | <.001 |

Note: DyNAMiC: 525 eligible observations from 177 participants. Linear mixed model with participant-specific intercepts; rest and male sex are the reference categories. Age is mean centered within the cohort and divided by the cohort SD, with Age² its square. *p*-values are two-sided and unadjusted.

**Table S3. Sensitivity to weak structural connections**

**A. Global SFC state contrasts**

| **Cohort** | **Weak edges removed** | **Contrast** | ***Δ* [95% CI]** | ***p*** |
| --- | --- | --- | --- | --- |
| DyNAMiC | 0% | Movie-Rest | 0.0090 [0.0046, 0.0134] | <.001 |
| DyNAMiC | 0% | *N*-back-Rest | −0.0127 [−0.0171, −0.0084] | <.001 |
| DyNAMiC | 25% | Movie-Rest | 0.0101 [0.0055, 0.0146] | <.001 |
| DyNAMiC | 25% | *N*-back-Rest | −0.0134 [−0.0180, −0.0089] | <.001 |
| DyNAMiC | 50% | Movie-Rest | 0.0114 [0.0066, 0.0162] | <.001 |
| DyNAMiC | 50% | *N*-back-Rest | −0.0147 [−0.0195, −0.0100] | <.001 |
| CamCAN | 0% | Movie-Rest | 0.0130 [0.0094, 0.0167] | <.001 |
| CamCAN | 0% | SMT-Rest | 0.0105 [0.0072, 0.0137] | <.001 |
| CamCAN | 25% | Movie-Rest | 0.0182 [0.0141, 0.0223] | <.001 |
| CamCAN | 25% | SMT-Rest | 0.0127 [0.0089, 0.0164] | <.001 |
| CamCAN | 50% | Movie-Rest | 0.0221 [0.0176, 0.0265] | <.001 |
| CamCAN | 50% | SMT-Rest | 0.0140 [0.0100, 0.0180] | <.001 |

**B. DyNAMiC connectivity-ratio contrasts**

| **Weak edges removed** | **Contrast** | ***Δ* [95% CI]** | ***p*** |
| --- | --- | --- | --- |
| 0% | Movie-Rest | 0.0111 [−0.0074, 0.0297] | 0.239 |
| 0% | *N*-back-Rest | −0.0283 [−0.0467, −0.0098] | 0.003 |
| 0% | Movie-*N*-back | 0.0394 [0.0208, 0.0580] | <.001 |
| 25% | Movie-Rest | 0.0080 [−0.0100, 0.0259] | 0.384 |
| 25% | *N*-back-Rest | −0.0348 [−0.0527, −0.0170] | <.001 |
| 25% | Movie-*N*-back | 0.0428 [0.0248, 0.0608] | <.001 |
| 50% | Movie-Rest | 0.0060 [−0.0120, 0.0240] | 0.511 |
| 50% | *N*-back-Rest | −0.0393 [−0.0572, −0.0214] | <.001 |
| 50% | Movie-*N*-back | 0.0453 [0.0273, 0.0633] | <.001 |

Note: Thresholds removed the weakest 25% or 50% of SC weights within each participant, with the same edge mask applied across that participant’s states. The original state model was retained. The connectivity-ratio reference remained edges with originally zero SC; weak edges were omitted, not reassigned to the reference set. *p*-values are two-sided and unadjusted.

**Table S4. CamCAN primary state model**

| **Predictor** | **Estimate** | **SE** | **95% CI** | ***t*** | ***df*** | ***p*** |
| --- | --- | --- | --- | --- | --- | --- |
| Intercept | 0.2834 | 0.0037 | [0.2761, 0.2907] | 75.87 | 1147 | <.001 |
| Movie vs. rest | 0.0130 | 0.0019 | [0.0094, 0.0167] | 7.02 | 1147 | <.001 |
| Sensorimotor vs. rest | 0.0105 | 0.0017 | [0.0072, 0.0137] | 6.26 | 1147 | <.001 |
| Age (centered) | -0.0002 | 6.36e-05 | [-0.0003, -9.40e-05] | -3.44 | 1147 | <.001 |
| Sex (male vs. female) | 0.0011 | 0.0022 | [-0.0031, 0.0054] | 0.53 | 1147 | .593 |
| Mean FD | -0.0551 | 0.0172 | [-0.0888, -0.0215] | -3.21 | 1147 | .001 |

Note: CamCAN: 1153 observations from 546 participants. Linear mixed model with participant-specific intercepts; rest and female sex are the reference categories. Age is mean centered within the cohort and divided by the cohort SD, with Age² its square. Omnibus state test: *F*(2,1147) = 27.90, *p* < .001. *p*-values are two-sided and unadjusted.

**Table S5. DyNAMiC linear working-memory load model**

| **Predictor** | **Estimate** | **SE** | **95% CI** | ***t*** | ***df*** | ***p*** |
| --- | --- | --- | --- | --- | --- | --- |
| Intercept | 0.2020 | 0.0037 | [0.1947, 0.2093] | 54.26 | 519 | <.001 |
| Age | −0.0104 | 0.0021 | [−0.0146, −0.0062] | −4.85 | 519 | <.001 |
| Sex (female vs. male) | 0.0048 | 0.0041 | [−0.0033, 0.0129] | 1.16 | 519 | .245 |
| Mean FD | −0.0091 | 0.0015 | [−0.0120, −0.0062] | −6.17 | 519 | <.001 |
| Age² | 0.0066 | 0.0024 | [0.0018, 0.0113] | 2.71 | 519 | .007 |
| Linear load | −0.0022 | 0.0007 | [−0.0035, −0.0009] | −3.30 | 519 | .001 |

Note: 525 load observations from 175 participants. Load is coded −1, 0, 1 for 1-back, 2-back, 3-back, respectively. Eligibility follows the full parent n-back run. *p*-values are two-sided and unadjusted.

**Table S6. System segregation**

| **Predictor** | **Estimate** | **SE** | **95% CI** | ***t*** | ***df*** | ***p*** |
| --- | --- | --- | --- | --- | --- | --- |
| Intercept | 0.4227 | 0.0162 | [0.3908, 0.4546] | 26.05 | 518 | <.001 |
| Movie vs. rest | −0.0371 | 0.0088 | [−0.0545, −0.0197] | −4.20 | 518 | <.001 |
| N-back vs. rest | −0.1395 | 0.0088 | [−0.1568, −0.1223] | −15.89 | 518 | <.001 |
| Age | −0.0026 | 0.0089 | [−0.0201, 0.0149] | −0.29 | 518 | .769 |
| Sex (female vs. male) | 0.0543 | 0.0171 | [0.0207, 0.0879] | 3.17 | 518 | .002 |
| Mean FD | −0.0308 | 0.0063 | [−0.0431, −0.0185] | −4.92 | 518 | <.001 |
| Age² | 0.0257 | 0.0099 | [0.0062, 0.0452] | 2.58 | 518 | .010 |

Note: DyNAMiC: 525 observations from 177 participants. Linear mixed model with participant-specific intercepts; rest and male sex are the reference categories. Age is mean centered within the cohort and divided by the cohort SD, with Age² its square. Omnibus state test: *F*(2,518) = 135.19, *p* < .001. *p*-values are two-sided and unadjusted.

**Table S7. DyNAMiC system-segregation-adjusted SFC**

| **Predictor** | **Estimate** | **SE** | **95% CI** | ***t*** | ***df*** | ***p*** |
| --- | --- | --- | --- | --- | --- | --- |
| Intercept | 0.2408 | 0.0036 | [0.2337, 0.2479] | 66.85 | 517 | <.001 |
| Movie vs. rest | 0.0115 | 0.0021 | [0.0073, 0.0157] | 5.39 | 517 | <.001 |
| N-back vs. rest | 0.0033 | 0.0025 | [−0.0017, 0.0082] | 1.31 | 517 | .191 |
| Age | −0.0065 | 0.0020 | [−0.0104, −0.0026] | −3.29 | 517 | .001 |
| Sex (female vs. male) | 0.0015 | 0.0038 | [−0.0060, 0.0090] | 0.39 | 517 | .694 |
| Mean FD | −0.0136 | 0.0015 | [−0.0165, −0.0107] | −9.21 | 517 | <.001 |
| Age² | 0.0035 | 0.0022 | [−0.0009, 0.0078] | 1.57 | 517 | .116 |
| System segregation | 0.0167 | 0.0015 | [0.0137, 0.0197] | 10.97 | 517 | <.001 |

Note: Same observations and covariates as Table S6. The matched base model gives *β* = 0.0074, 95% CI [0.0030, 0.0117], *t*(518) = 3.31, *p* < 0.001 for Movie−Rest and *β* = −0.0122, 95% CI [−0.0165, −0.0079], *t*(518) = −5.52, *p* < 0.001 for N-back−Rest. *p*-values are two-sided and unadjusted.

**Table S7 continued. State-Segregation interactions**

| **Predictor** | **Estimate** | **SE** | **95% CI** | ***t*** | ***df*** | ***p*** |
| --- | --- | --- | --- | --- | --- | --- |
| Intercept | 0.2422 | 0.0036 | [0.2352, 0.2492] | 67.69 | 515 | <.001 |
| Movie vs. rest | 0.0102 | 0.0022 | [0.0059, 0.0145] | 4.65 | 515 | <.001 |
| N-back vs. rest | 0.0054 | 0.0026 | [0.0002, 0.0106] | 2.05 | 515 | .041 |
| Age | −0.0062 | 0.0019 | [−0.0100, −0.0024] | −3.21 | 515 | .001 |
| Sex (female vs. male) | 0.0017 | 0.0037 | [−0.0057, 0.0090] | 0.44 | 515 | .657 |
| Mean FD | −0.0138 | 0.0015 | [−0.0166, −0.0109] | −9.39 | 515 | <.001 |
| Age² | 0.0031 | 0.0022 | [−0.0011, 0.0074] | 1.45 | 515 | .148 |
| System segregation | 0.0135 | 0.0018 | [0.0099, 0.0172] | 7.34 | 515 | <.001 |
| Movie vs. rest × System segregation | 0.0041 | 0.0022 | [−0.0003, 0.0085] | 1.85 | 515 | .065 |
| N-back vs. rest × System segregation | 0.0093 | 0.0025 | [0.0043, 0.0143] | 3.65 | 515 | <.001 |

Note: Same 525 observations and covariates as the additive model, with the state-segregation interaction added: N-back−Rest, *β* = 0.0093, *t*(515) = 3.65, *p* < .001; N-back−Movie, *β* = 0.0051, *t*(515) = 2.02, *p* = .044; Movie−Rest, *β* = 0.0041, *t*(515) = 1.85, *p* = .065. *p*-values are two-sided and unadjusted.

**Table S8. Spatial correspondence and spherical-rotation tests**

| **Reference** | **State** | ***ρ*** | **Spin *p*** | ***p*_adj_** |
| --- | --- | --- | --- | --- |
| Gradient | Rest | 0.2557 | .087 | .087 |
| Gradient | Movie | 0.3585 | .018 | .053 |
| Gradient | *N*-back | 0.2686 | .087 | .087 |
| Group stability | Rest | 0.3719 | .001 | .002 |
| Group stability | Movie | 0.4278 | <.001 | .002 |
| Group stability | *N*-back | 0.3697 | .002 | .002 |

Note: *p*-values (*p*_adj_) are FDR-adjusted across the three state associations for each reference map, following 10,000 spins. Group maps use eligible runs in each state. Reference-map bootstraps are conditioned on the fitted gradient or group stability map.

**Table S9. DyNAMiC network state contrasts**

| **Network** | **Estimate** | **SE** | **95% CI** | ***t*** | ***df*** | ***p*_adj_** |
| --- | --- | --- | --- | --- | --- | --- |
| **Movie−Rest** | | | | | | |
| Visual1 | 0.0217 | 0.0038 | [0.0142, 0.0292] | 5.70 | 518 | <.001 |
| Visual2 | 0.0311 | 0.0030 | [0.0251, 0.0370] | 10.28 | 518 | <.001 |
| Somatomotor | −0.0064 | 0.0032 | [−0.0127, −3.96e−05] | −1.98 | 518 | .067 |
| Cingulo-Opercular | 0.0038 | 0.0028 | [−0.0017, 0.0093] | 1.35 | 518 | .200 |
| Dorsal-Attention | 0.0158 | 0.0026 | [0.0107, 0.0210] | 6.00 | 518 | <.001 |
| Language | 0.0263 | 0.0031 | [0.0202, 0.0323] | 8.53 | 518 | <.001 |
| Frontoparietal | −0.0120 | 0.0023 | [−0.0166, −0.0075] | −5.17 | 518 | <.001 |
| Auditory | 0.0384 | 0.0046 | [0.0293, 0.0474] | 8.32 | 518 | <.001 |
| Posterior-Multimodal | 0.0219 | 0.0033 | [0.0154, 0.0285] | 6.58 | 518 | <.001 |
| Default | −0.0151 | 0.0025 | [−0.0199, −0.0102] | −6.12 | 518 | <.001 |
| Orbito-Affective | 0.0031 | 0.0050 | [−0.0068, 0.0130] | 0.61 | 518 | .589 |
| Ventral-Multimodal | 0.0120 | 0.0065 | [−0.0006, 0.0247] | 1.87 | 518 | .084 |
| **N-back−Rest** | | | | | | |
| Visual1 | 0.0227 | 0.0038 | [0.0152, 0.0301] | 5.99 | 518 | <.001 |
| Visual2 | −0.0055 | 0.0030 | [−0.0114, 0.0004] | −1.83 | 518 | .087 |
| Somatomotor | −0.0256 | 0.0032 | [−0.0319, −0.0193] | −8.01 | 518 | <.001 |
| Cingulo-Opercular | −0.0047 | 0.0028 | [−0.0102, 0.0008] | −1.67 | 518 | .119 |
| Dorsal-Attention | 0.0145 | 0.0026 | [0.0094, 0.0197] | 5.54 | 518 | <.001 |
| Language | −0.0168 | 0.0031 | [−0.0229, −0.0108] | −5.50 | 518 | <.001 |
| Frontoparietal | −0.0067 | 0.0023 | [−0.0112, −0.0021] | −2.89 | 518 | .006 |
| Auditory | −0.0279 | 0.0046 | [−0.0369, −0.0189] | −6.10 | 518 | <.001 |
| Posterior-Multimodal | −0.0072 | 0.0033 | [−0.0137, −0.0007] | −2.17 | 518 | .044 |
| Default | −0.0186 | 0.0024 | [−0.0234, −0.0138] | −7.60 | 518 | <.001 |
| Orbito-Affective | −0.0146 | 0.0050 | [−0.0244, −0.0047] | −2.91 | 518 | .006 |
| Ventral-Multimodal | 0.0097 | 0.0064 | [−0.0030, 0.0223] | 1.50 | 518 | .160 |
| **Movie−N-back** | | | | | | |
| Visual1 | −0.0010 | 0.0038 | [−0.0085, 0.0065] | −0.25 | 518 | .799 |
| Visual2 | 0.0366 | 0.0030 | [0.0306, 0.0425] | 12.09 | 518 | <.001 |
| Somatomotor | 0.0193 | 0.0032 | [0.0129, 0.0256] | 5.98 | 518 | <.001 |
| Cingulo-Opercular | 0.0085 | 0.0028 | [0.0029, 0.0140] | 3.01 | 518 | .005 |
| Dorsal-Attention | 0.0013 | 0.0026 | [−0.0039, 0.0065] | 0.50 | 518 | .657 |
| Language | 0.0431 | 0.0031 | [0.0371, 0.0492] | 13.98 | 518 | <.001 |
| Frontoparietal | −0.0054 | 0.0023 | [−0.0099, −0.0008] | −2.30 | 518 | .033 |
| Auditory | 0.0663 | 0.0046 | [0.0572, 0.0754] | 14.38 | 518 | <.001 |
| Posterior-Multimodal | 0.0291 | 0.0033 | [0.0226, 0.0357] | 8.73 | 518 | <.001 |
| Default | 0.0035 | 0.0025 | [−0.0013, 0.0084] | 1.43 | 518 | .177 |
| Orbito-Affective | 0.0177 | 0.0050 | [0.0078, 0.0276] | 3.51 | 518 | <.001 |
| Ventral-Multimodal | 0.0024 | 0.0065 | [−0.0103, 0.0151] | 0.37 | 518 | .732 |

Note: Participant-level network means are analyzed using linear mixed models of state contrasts with participant-specific intercepts, age, sex, and mean FD as covariates. *p*-values (*p*_adj_) are FDR-adjusted across 36 contrasts; 95% confidence intervals are unadjusted.

**Table S10. CamCAN network state contrasts**

| **Network** | **Estimate** | **SE** | **95% CI** | ***t*** | ***df*** | ***p*_adj_** |
| --- | --- | --- | --- | --- | --- | --- |
| **Movie−Rest** | | | | | | |
| Visual1 | 0.0156 | 0.0053 | [0.0051, 0.0261] | 2.92 | 1147 | 0.004 |
| Visual2 | 0.0091 | 0.0029 | [0.0033, 0.0149] | 3.08 | 1147 | 0.003 |
| Somatomotor | −0.0157 | 0.0033 | [−0.0221, −0.0092] | −4.79 | 1147 | <.001 |
| Cingulo−Opercular | −0.0091 | 0.0024 | [−0.0138, −0.0043] | −3.72 | 1147 | <.001 |
| Dorsal−Attention | 0.0197 | 0.0022 | [0.0153, 0.0241] | 8.76 | 1147 | <.001 |
| Language | 0.0321 | 0.0029 | [0.0265, 0.0377] | 11.23 | 1147 | <.001 |
| Frontoparietal | 0.0110 | 0.0026 | [0.0059, 0.0161] | 4.24 | 1147 | <.001 |
| Auditory | −0.0037 | 0.0035 | [−0.0107, 0.0032] | −1.06 | 1147 | 0.287 |
| Posterior−Multimodal | 0.0173 | 0.0031 | [0.0113, 0.0233] | 5.65 | 1147 | <.001 |
| Default | 0.0424 | 0.0024 | [0.0377, 0.0470] | 18.01 | 1147 | <.001 |
| Orbito−Affective | 0.1703 | 0.0062 | [0.1581, 0.1826] | 27.27 | 1147 | <.001 |
| Ventral−Multimodal | 0.0779 | 0.0057 | [0.0666, 0.0891] | 13.55 | 1147 | <.001 |
| **Sensorimotor−Rest** | | | | | | |
| Visual1 | 0.0251 | 0.0050 | [0.0152, 0.0350] | 4.98 | 1147 | <.001 |
| Visual2 | 0.0098 | 0.0026 | [0.0047, 0.0149] | 3.78 | 1147 | <.001 |
| Somatomotor | 0.0275 | 0.0030 | [0.0216, 0.0333] | 9.22 | 1147 | <.001 |
| Cingulo−Opercular | −0.0132 | 0.0022 | [−0.0176, −0.0088] | −5.93 | 1147 | <.001 |
| Dorsal−Attention | 0.0222 | 0.0020 | [0.0183, 0.0261] | 11.18 | 1147 | <.001 |
| Language | 0.0148 | 0.0026 | [0.0096, 0.0200] | 5.60 | 1147 | <.001 |
| Frontoparietal | 0.0053 | 0.0024 | [0.0006, 0.0101] | 2.21 | 1147 | 0.030 |
| Auditory | 0.0040 | 0.0031 | [−0.0020, 0.0100] | 1.31 | 1147 | 0.192 |
| Posterior−Multimodal | 0.0063 | 0.0028 | [0.0009, 0.0117] | 2.28 | 1147 | 0.029 |
| Default | 0.0097 | 0.0022 | [0.0053, 0.0140] | 4.39 | 1147 | <.001 |
| Orbito−Affective | 0.0133 | 0.0059 | [0.0017, 0.0249] | 2.25 | 1147 | 0.029 |
| Ventral−Multimodal | 0.0283 | 0.0054 | [0.0178, 0.0388] | 5.28 | 1147 | <.001 |

Note: Participant-level network means are analyzed using linear mixed models of state contrasts with participant-specific intercepts, age, sex, and mean FD as covariates. *p*-values (*p*_adj_) are FDR-adjusted across 12 tests within each contrast; 95% confidence intervals are unadjusted.

**Table S11. Functional reconfiguration and age**

| **Predictor** | **Estimate** | **SE** | **95% CI** | ***t*** | ***df*** | ***p*** |
| --- | --- | --- | --- | --- | --- | --- |
| Intercept | −0.0021 | 0.0085 | [−0.0189, 0.0146] | −0.25 | 338 | .802 |
| N-back vs. movie | −0.0358 | 0.0087 | [−0.0529, −0.0187] | −4.11 | 338 | <.001 |
| Age | 0.0021 | 0.0005 | [0.0012, 0.0030] | 4.70 | 338 | <.001 |
| Sex (female vs. male) | −0.0088 | 0.0042 | [−0.0171, −0.0006] | −2.11 | 338 | .036 |
| Mean FD | −0.0064 | 0.0021 | [−0.0107, −0.0022] | −3.01 | 338 | .003 |
| Reconfiguration | 0.0758 | 0.0225 | [0.0317, 0.1200] | 3.38 | 338 | <.001 |
| N-back vs. movie × Age | −0.0021 | 0.0005 | [−0.0030, −0.0011] | −4.15 | 338 | <.001 |
| N-back vs. movie × Reconfiguration | 0.0146 | 0.0209 | [−0.0266, 0.0557] | 0.70 | 338 | .486 |
| Age × Reconfiguration | −0.0060 | 0.0011 | [−0.0082, −0.0038] | −5.32 | 338 | <.001 |
| N-back vs. movie × Age × Reconfiguration | 0.0058 | 0.0012 | [0.0035, 0.0082] | 4.95 | 338 | <.001 |

Note: 348 eligible task–rest observations. Linear mixed model with participant-specific intercepts. The outcome is task-minus-rest SFC. Movie is the reference task. The direct three-way interaction tests whether age-related reconfiguration slopes differ between tasks. The derived age × reconfiguration effect during N-back was not significant (*t*(338) = −0.15, *p* = .883). *p*-values are two-sided and unadjusted.

**Table S12. Global SFC age associations**

| **Cohort** | **State** | **Age estimate [95% CI]** | **Age *p*_adj_** | **Age^2^ estimate [95% CI]** | **Age^2^ *p*_adj_** | **Overall age *p*_adj_** |
| --- | --- | --- | --- | --- | --- | --- |
| DyNAMiC | Rest | -0.0041 [-0.0072, -0.0010] | 0.013 | 0.00128 [-0.00070, 0.00325] | 0.204 | 0.015 |
| DyNAMiC | Movie | -0.0055 [-0.0086, -0.0024] | 0.002 | 0.00173 [-0.00025, 0.00371] | 0.131 | 0.002 |
| DyNAMiC | *N*-back | -0.0030 [-0.0060, 0.0001] | 0.059 | 0.00324 [0.00126, 0.00523] | 0.004 | 0.002 |
| CamCAN | Rest | -0.0004 [-0.0022, 0.0015] | 0.694 | 0.00237 [0.00138, 0.00336] | <.001 | <.001 |
| CamCAN | Movie | -0.0027 [-0.0041, -0.0013] | <.001 | 0.00136 [0.00060, 0.00212] | <.001 | <.001 |
| CamCAN | SMT | -0.0026 [-0.0042, -0.0009] | 0.003 | 0.00185 [0.00092, 0.00278] | <.001 | <.001 |

Note: Age is mean centered within each cohort and divided by 10. Linear estimates are from linear-only mixed models, in SFC *r* units per decade; quadratic estimates are squared-age coefficients from quadratic mixed models, in *r* units per decade squared. Models include sex, the cohort-specific mean FD covariates, and participant-specific intercepts. 95% confidence intervals are unadjusted. *p*-values (*p*_adj_) are FDR-adjusted across the three states within each cohort, separately for linear, quadratic, and overall age tests. The overall age test jointly assesses the linear and quadratic terms.

**Table S13. State differences in global SFC age associations**

| **Cohort** | **State pair** | **Slope difference [95% CI]** | **Age Linear *p*_adj_** | **Overall age *p*_adj_** |
| --- | --- | --- | --- | --- |
| DyNAMiC | Movie - Rest | -0.0014 [-0.0039, 0.0012] | 0.368 | 0.493 |
| DyNAMiC | *N*-back - Rest | 0.0012 [-0.0014, 0.0037] | 0.368 | 0.072 |
| DyNAMiC | *N*-back - Movie | 0.0025 [-0.0000, 0.0051] | 0.154 | 0.072 |
| CamCAN | Movie - Rest | -0.0023 [-0.0041, -0.0005] | 0.032 | 0.008 |
| CamCAN | SMT - Rest | -0.0022 [-0.0041, -0.0003] | 0.032 | 0.066 |
| CamCAN | SMT - Movie | 0.0001 [-0.0015, 0.0017] | 0.907 | 0.537 |

Note: State comparisons are first state minus second state. Slope differences are from linear-only mixed models, in SFC *r* units per decade, with unadjusted 95% confidence intervals. Age is mean centered within each cohort and divided by 10. Primary models include sex, mean FD, and participant-specific intercepts. The linear and overall age columns report primary-model tests. Overall age tests assess between-state differences in the linear and quadratic age coefficients together. Pairwise *p*-values (*p*_adj_) are FDR-adjusted across the three state pairs within each cohort, separately for linear and overall age tests. Overall state-by-age interactions: DyNAMiC, *F*(4, 514) = 2.39, *p* = .0497; CamCAN, *F*(4, 1142) = 3.17, *p* = .013.

**Table S14. DyNAMiC network-level age associations**

| **Network** | **Linear age slope [95% CI]** | **Linear age *p*_adj_** | **Quadratic estimate** | **Quadratic *p*_adj_** | **Joint age *p*_adj_** |
| --- | --- | --- | --- | --- | --- |
| **Rest** | | | | | |
| **Visual1** | -0.0075 [-0.0121, -0.0029] | 0.005 | 0.00296 | 0.119 | **0.003** |
| Visual2 | -0.0011 [-0.0054, 0.0032] | 0.661 | 0.00316 | 0.088 | 0.096 |
| **Somatomotor** | -0.0064 [-0.0104, -0.0024] | 0.005 | 0.00261 | 0.118 | **0.003** |
| **Cingulo-Opercular** | -0.0060 [-0.0095, -0.0024] | 0.004 | 0.00110 | 0.414 | **0.007** |
| **Dorsal-Attention** | 0.0033 [-0.0002, 0.0069] | 0.119 | 0.00258 | 0.088 | **0.026** |
| Language | -0.0049 [-0.0091, -0.0007] | 0.054 | 0.00036 | 0.819 | 0.103 |
| Frontoparietal | -0.0008 [-0.0038, 0.0023] | 0.661 | 0.00158 | 0.181 | 0.297 |
| **Auditory** | -0.0238 [-0.0300, -0.0176] | <.001 | -0.00363 | 0.141 | **<.001** |
| Posterior-Multimodal | -0.0021 [-0.0063, 0.0022] | 0.471 | -0.00064 | 0.725 | 0.647 |
| Default | -0.0028 [-0.0061, 0.0006] | 0.177 | 0.00159 | 0.208 | 0.126 |
| **Orbito-Affective** | -0.0108 [-0.0169, -0.0047] | 0.003 | 0.00362 | 0.141 | **0.002** |
| Ventral-Multimodal | -0.0026 [-0.0100, 0.0048] | 0.595 | 0.00144 | 0.642 | 0.718 |
| **Movie** | | | | | |
| **Visual1** | -0.0073 [-0.0120, -0.0027] | 0.006 | 0.00362 | 0.087 | **0.002** |
| Visual2 | -0.0039 [-0.0083, 0.0004] | 0.131 | 0.00225 | 0.181 | 0.088 |
| **Somatomotor** | -0.0092 [-0.0132, -0.0051] | <.001 | 0.00239 | 0.141 | **<.001** |
| **Cingulo-Opercular** | -0.0074 [-0.0110, -0.0039] | <.001 | 0.00169 | 0.208 | **<.001** |
| Dorsal-Attention | 0.0018 [-0.0018, 0.0054] | 0.471 | 0.00244 | 0.112 | 0.095 |
| **Language** | -0.0062 [-0.0105, -0.0019] | 0.012 | 0.00061 | 0.725 | **0.030** |
| Frontoparietal | -0.0013 [-0.0043, 0.0018] | 0.551 | 0.00220 | 0.088 | 0.089 |
| **Auditory** | -0.0218 [-0.0280, -0.0156] | <.001 | -0.00274 | 0.230 | **<.001** |
| Posterior-Multimodal | -0.0004 [-0.0047, 0.0039] | 0.879 | -0.00021 | 0.883 | 0.979 |
| **Default** | -0.0065 [-0.0099, -0.0031] | <.001 | 0.00252 | 0.088 | **<.001** |
| **Orbito-Affective** | -0.0124 [-0.0186, -0.0063] | <.001 | 0.00323 | 0.181 | **<.001** |
| Ventral-Multimodal | 0.0019 [-0.0056, 0.0094] | 0.661 | 0.00082 | 0.783 | 0.850 |
| **N-back** | | | | | |
| Visual1 | -0.0037 [-0.0083, 0.0009] | 0.182 | 0.00307 | 0.118 | 0.057 |
| **Visual2** | -0.0015 [-0.0058, 0.0028] | 0.595 | 0.00456 | 0.009 | **0.009** |
| **Somatomotor** | -0.0070 [-0.0111, -0.0030] | 0.003 | 0.00461 | 0.009 | **<.001** |
| **Cingulo-Opercular** | -0.0041 [-0.0077, -0.0006] | 0.053 | 0.00222 | 0.131 | **0.023** |
| **Dorsal-Attention** | 0.0020 [-0.0016, 0.0056] | 0.415 | 0.00393 | 0.009 | **0.005** |
| Language | -0.0043 [-0.0086, -0.0001] | 0.091 | 0.00192 | 0.230 | 0.087 |
| **Frontoparietal** | 0.0012 [-0.0019, 0.0042] | 0.576 | 0.00334 | 0.009 | **0.006** |
| **Auditory** | -0.0201 [-0.0263, -0.0140] | <.001 | -0.00212 | 0.371 | **<.001** |
| Posterior-Multimodal | -0.0013 [-0.0055, 0.0030] | 0.653 | 0.00214 | 0.205 | 0.320 |
| **Default** | -0.0036 [-0.0070, -0.0002] | 0.078 | 0.00364 | 0.009 | **0.002** |
| **Orbito-Affective** | -0.0057 [-0.0119, 0.0004] | 0.119 | 0.00543 | 0.042 | **0.009** |
| Ventral-Multimodal | -0.0004 [-0.0078, 0.0071] | 0.926 | 0.00365 | 0.205 | 0.370 |

Note: Linear mixed models of age associations include participant-specific intercepts, sex, and mean FD as covariates. *p*-values (*p*_adj_) are FDR-adjusted across 12 networks × 3 states within each cohort, separately for linear, quadratic, and joint age tests. Bold indicates significant joint age associations.

**Table S15. CamCAN network-level age associations**

| **Network** | **Linear age slope [95% CI]** | **Linear age *p*_adj_** | **Quadratic estimate** | **Quadratic *p*_adj_** | **Joint age *p*_adj_** |
| --- | --- | --- | --- | --- | --- |
| **Rest** | | | | | |
| **Visual1** | -0.0097 [-0.0146, -0.0048] | <.001 | -0.00085 | 0.587 | **<.001** |
| **Visual2** | -0.0041 [-0.0071, -0.0012] | 0.009 | 0.00352 | <.001 | **<.001** |
| **Somatomotor** | -0.0034 [-0.0065, -0.0003] | 0.041 | 0.00189 | 0.053 | **0.010** |
| **Cingulo-Opercular** | -0.0015 [-0.0038, 0.0008] | 0.275 | 0.00275 | <.001 | **<.001** |
| **Dorsal-Attention** | 0.0050 [0.0027, 0.0072] | <.001 | 0.00392 | <.001 | **<.001** |
| **Language** | 0.0031 [0.0004, 0.0058] | 0.036 | 0.00180 | 0.032 | **0.006** |
| **Frontoparietal** | -0.0003 [-0.0027, 0.0022] | 0.862 | 0.00276 | <.001 | **<.001** |
| **Auditory** | -0.0057 [-0.0094, -0.0021] | 0.004 | -0.00179 | 0.126 | **0.002** |
| **Posterior-Multimodal** | 0.0015 [-0.0014, 0.0045] | 0.390 | 0.00284 | 0.002 | **0.003** |
| **Default** | -0.0040 [-0.0062, -0.0018] | <.001 | 0.00192 | 0.005 | **<.001** |
| **Orbito-Affective** | -0.0130 [-0.0188, -0.0072] | <.001 | 0.00298 | 0.112 | **<.001** |
| **Ventral-Multimodal** | -0.0066 [-0.0120, -0.0012] | 0.025 | 0.00736 | <.001 | **<.001** |
| **Movie** | | | | | |
| **Visual1** | -0.0061 [-0.0098, -0.0024] | 0.003 | 0.00146 | 0.200 | **0.002** |
| **Visual2** | 0.0007 [-0.0017, 0.0030] | 0.636 | 0.00395 | <.001 | **<.001** |
| **Somatomotor** | -0.0144 [-0.0168, -0.0120] | <.001 | 0.00019 | 0.788 | **<.001** |
| **Cingulo-Opercular** | -0.0056 [-0.0073, -0.0038] | <.001 | 0.00119 | 0.029 | **<.001** |
| **Dorsal-Attention** | 0.0043 [0.0025, 0.0060] | <.001 | 0.00250 | <.001 | **<.001** |
| Language | -0.0006 [-0.0027, 0.0014] | 0.636 | 0.00017 | 0.788 | 0.795 |
| Frontoparietal | -0.0019 [-0.0038, -0.0001] | 0.053 | 0.00063 | 0.270 | 0.052 |
| **Auditory** | -0.0060 [-0.0089, -0.0030] | <.001 | 0.00022 | 0.788 | **<.001** |
| **Posterior-Multimodal** | 0.0031 [0.0008, 0.0054] | 0.014 | 0.00201 | 0.004 | **<.001** |
| **Default** | -0.0041 [-0.0057, -0.0024] | <.001 | 0.00069 | 0.172 | **<.001** |
| **Orbito-Affective** | -0.0137 [-0.0180, -0.0094] | <.001 | -0.00106 | 0.422 | **<.001** |
| **Ventral-Multimodal** | 0.0000 [-0.0040, 0.0041] | 0.985 | 0.00367 | 0.003 | **0.005** |
| **SMT** | | | | | |
| **Visual1** | -0.0167 [-0.0211, -0.0122] | <.001 | 0.00216 | 0.134 | **<.001** |
| **Visual2** | -0.0083 [-0.0110, -0.0056] | <.001 | 0.00339 | <.001 | **<.001** |
| **Somatomotor** | -0.0119 [-0.0147, -0.0090] | <.001 | 0.00140 | 0.128 | **<.001** |
| **Cingulo-Opercular** | 0.0008 [-0.0013, 0.0029] | 0.569 | 0.00249 | <.001 | **<.001** |
| **Dorsal-Attention** | 0.0034 [0.0013, 0.0054] | 0.003 | 0.00346 | <.001 | **<.001** |
| **Language** | -0.0028 [-0.0053, -0.0004] | 0.033 | 0.00084 | 0.278 | **0.035** |
| **Frontoparietal** | 0.0004 [-0.0018, 0.0026] | 0.764 | 0.00178 | 0.012 | **0.021** |
| **Auditory** | -0.0096 [-0.0130, -0.0062] | <.001 | -0.00092 | 0.399 | **<.001** |
| **Posterior-Multimodal** | 0.0005 [-0.0022, 0.0032] | 0.764 | 0.00233 | 0.006 | **0.012** |
| **Default** | -0.0044 [-0.0064, -0.0024] | <.001 | 0.00094 | 0.139 | **<.001** |
| **Orbito-Affective** | -0.0079 [-0.0131, -0.0028] | 0.005 | 0.00264 | 0.126 | **0.003** |
| **Ventral-Multimodal** | -0.0074 [-0.0123, -0.0026] | 0.005 | 0.00353 | 0.024 | **<.001** |

Note: Linear mixed models of age associations include participant-specific intercepts, sex, and mean FD as covariates. *p*-values (*p*_adj_) are FDR-adjusted across 12 networks × 3 states within each cohort, separately for linear, quadratic, and joint age tests. Bold indicates significant joint age associations.
